# Sympathetic axonal sprouting induces changes in macrophage populations and protects against pancreatic cancer

**DOI:** 10.1101/2020.04.27.061648

**Authors:** Jérémy Guillot, Chloé Dominici, Adrien Lucchesi, Thi Trang Huyen Nguyen, Jérémy Nigri, Fabienne Guillaumond, Martin Bigonnet, Nelson Dusetti, Anders Etzerodt, Toby Lawrence, Pierre Pudlo, Florence Hubert, Jean-Yves Scoazec, Serge A. van de Pavert, Richard Tomasini, Sophie Chauvet, Fanny Mann

## Abstract

Recent evidence has highlighted the presence of neuronal nerve processes in the tumor microenvironment. However, the origin of intra-tumoral nerves remains poorly known, in part because of technical difficulties in tracing nerve fibers via regular histological preparations. Here, we employed three-dimensional (3D) imaging of cleared tissues for a comprehensive analysis of sympathetic innervation in pancreatic ductal adenocarcinoma (PDAC). The results support two independent, but coexisting, mechanisms: passive engulfment of pre-existing nerves within tumors and active, localized sprouting of nerve terminals into non-neoplastic lesions and tumor periphery. Nerve ablation revealed an inverse correlation between sympathetic innervation and tumor growth and spread. Furthermore, sympathectomy increased CD163^+^ macrophage levels, which contributed to worse outcomes. Altogether, our findings revealed protective properties of the sympathetic nervous system in PDAC immunity and progression that could pave the way for new treatments.

## Introduction

The peripheral nervous system (PNS) includes a large network of nerves that relays information back and forth between the brain and the body. Typically, a peripheral nerve contains thousands of nerve fibers, or axons, wrapped in bundles that defasciculate into individual axons and form complex branched networks within their target peripheral organ. Axons from the autonomic division of the PNS innervate and regulate most internal organs, glands, and blood vessels to maintain body homeostasis under basal and stress conditions. The autonomic nervous system also plays crucial roles in the regeneration process that restores organ structure and function after damage, leading to the concept of “nerve dependence” in tissue regeneration ^1^. Recent insights have revealed a role for the autonomic nervous system in promoting tumorigenesis, which can sometimes be considered an uncontrolled tissue regeneration process ^2^. The autonomic nervous system is divided into the sympathetic and parasympathetic nervous systems, whose effects on cancers depend on the type and stage. In prostate cancer, experimental ablation of adrenergic sympathetic nerves inhibits tumor initiation, whereas blocking cholinergic activity of parasympathetic nerves inhibits metastatic spreading at advanced stages of the disease ^3^. Similar pro-tumor activity of the autonomic nervous system has been reported in gastric cancer, where parasympathetic nerve fibers play a main role in promoting tumor initiation and progression ^4^. This has led to the notion that many cancers depend on nerves for development.

Pancreatic ductal adenocarcinoma (PDAC) is a severe cancer with poor prognosis that develops from the exocrine part of the pancreas. Both parasympathetic and sympathetic nervous systems densely innervate the pancreas and regulate exocrine secretions of acinar and ductal cells ^5^. The role of the autonomic nervous system in PDAC development has been addressed in recent studies that have revealed a more complex and unexpected picture than that suggested by research on other cancer models. Indeed, the interruption of parasympathetic innervation or activity strongly promotes PDAC progression in *Kras*-mutated mouse models, revealing an unexpected anti-tumor effect of parasympathetic cholinergic signaling ^6,7^. The role of the sympathetic nervous system in PDAC has been studied in terms of tumor response to stress and has shown context-dependent effects. Experimental sympathectomy or blockade of β-adrenergic signaling in mouse models of PDAC abolishes the pro-tumor effect of chronic restraint stress, but also inhibits the anti-tumor effect of “eustress” (positive stress) induced by enriched housing conditions ^8–10^. How the reactivity of the sympathetic nervous system to stressors can have opposing actions on tumor development is not clear, but it may involve differential regulation of immune cell functions (eustress) or direct effects on tumor cell growth (distress). Thus far, the contribution of the sympathetic nervous system to PDAC development has not been explored in animals kept under basal unstressed conditions.

PNS control over tumorigenesis relies on its ability to innervate developing tumors, which are known to express large numbers of neurotrophic factors and axon guidance molecules ^11^. PDAC in particular is a cancer in which significant neuroplastic changes have long been described and which exhibits large nerve bundles in sections of resected tumors from human patients ^12,13^. Large nerves are also observed in murine PDAC tumors and an increased intra-tumoral density of sympathetic nerves has been reported in enlarged tumors of *Kras*-mutant mice subjected to chronic stress conditions ^8,14^. Stress-mediated upregulation of nerve growth factor (NGF) secretion by tumor cells was proposed to attract sympathetic nerve growth via “tumor axonogenesis” ^8^. However, a recent study proposed that axons innervating tumors may extend from new neurons generated by populations of precursor cells in the subventricular zone of the brain that would have to travel to the distant tumor through the bloodstream ^15^. Such a mechanism, initially described as contributing to the innervation of prostate cancer, has been proposed to occur in many other cancer types, including PDAC ^15^. Nonetheless, the relative contribution of axonogenesis and neurogenesis to neuroplastic changes accompanying the development and progression of PDAC remains to be explored.

In this study, we employed 3D imaging of optically cleared tissues to visualize sympathetic nerves and their terminal innervation patterns in pancreas and PDAC tumors. This approach allowed us to trace the origin of intra-tumoral nerves and accurately quantify changes in the morphology of axon networks innervating healthy or lesioned tissues, which are difficult to assess via regular 2D histological sections. In addition, we examined how sympathectomy of pancreatic tumors affects disease progression.

## Results

### Pre-existing sympathetic nerves become engulfed in pancreatic tumors

To study how the structure and distribution of sympathetic nerves change during PDAC development, we employed tyrosine hydroxylase (TH; an enzyme involved in biosynthesis of norepinephrine) antibody staining and optical tissue clearing to visualize adrenergic innervation of the whole murine pancreas by 3D light sheet fluorescent microscopy (LSFM). TH is expressed by postganglionic sympathetic fibers supplying the pancreas that originate from the coeliac-superior mesenteric complex. Although the enzyme is also present in some sensory neurons of the dorsal root ganglia (DRG) that innervate the skin and pelvic viscera, TH-expressing DRG neurons have not been reported in the pancreas ^16^. In 8-week-old mice, TH-labeled postganglionic sympathetic nerves entered the pancreas through a main entry point, then divided into two principal nerve fiber bundles that run in different directions and branch off at many points to innervate the whole organ (Fig. 1A, Supplementary Movie S1). One of these branches passed through the pancreas to innervate the adjacent spleen, which was kept attached to the pancreas during dissection (Fig. 1A).

**Figure 1.**
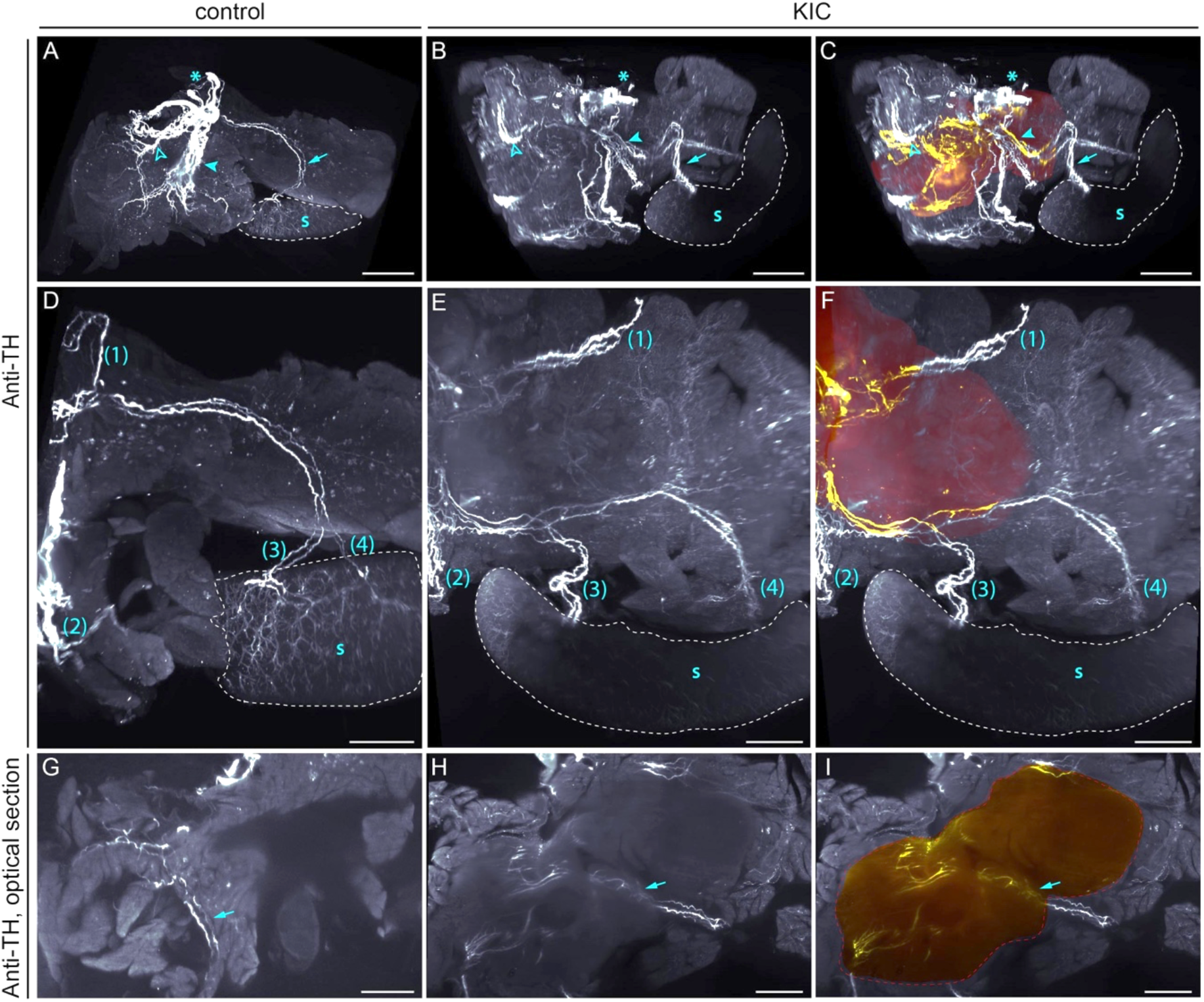
3D patterns of sympathetic nerve fiber bundles in control and KIC pancreata. Representative 3D views (**A–F**) and optical sections (**G–I**) of anti-TH labeled pancreas and spleen (s, delimited with dotted line) of 8-week-old control (**A, D**, and **G**) and KIC (**B**, **C**, **E**, **F**, **H**, and **I**) mice. All panels are LSFM images of solvent-cleared tissues. **A–C**, Asterisks indicate the nerve entry point into the pancreas; arrowheads indicate the main intra-pancreatic nerve tracks and the blue arrow points to the nerve bundle that innervates the spleen. **D–I**, Despite variations in their positioning, common nerve bundles (numbered from 1 to 4 in **D–F**, or indicated by the blue arrow in **G–I**) innervate the pancreas of control and KIC mice. **C, F, I**, 3D segmentation of pancreatic tumor (red), intra-tumoral TH^+^ nerve fibers (yellow), and extratumoral TH^+^ nerve fibers (white). Scale bars = 2 mm (**A**), 3 mm (**B, C**), and 1 mm (**D-I**).

We then compared this pattern of innervation to that of LSL-Kras^G12D/+^; Ink4a/Arf^lox/lox^; Pdx1-Cre (KIC) mice. By 8 weeks of age, all KIC mice have developed locally invasive PDAC ^17^. At this stage, the sympathetic nerve fiber bundles diverging from the main entry point could be recognized as in controls, but the overall pattern of pancreatic innervation appeared severely disrupted by tumor development (Fig. 1B, Supplementary Movie S2). To identify sympathetic nerves inside the tumor, we performed 3D segmentation of the PDAC, then traced and artificially colored the intra-tumoral nerve projections. Autofluorescence of the tissue (imaged at 488 nm excitation) provided sufficient structural information to accurately identify the contours of the tumor (Supplementary Fig. S1). We observed many bundles of sympathetic nerve fibers scattered in KIC tumors (Fig. 1C), confirming previous observations made in human samples and other genetically engineered mouse (GEM) models of PDAC ^8,18^. However, by tracing the whole course of the intra-tumoral nerves, we found that most corresponded to segments of en passant nerve bundles that were also present in the control pancreas (such as the nerve bundle projecting to the spleen; Fig. 1D–1I). These results suggest that the intra-tumoral sympathetic nerve bundles frequently reported in PDAC may correspond to pre-existing nerve bundles that pass through the pancreas and become embedded in the tumor as it develops.

### Remodeling of sympathetic axon nerve terminals

The above data suggest that the large sympathetic nerve bundles observed in pancreatic cancer are unlikely to result from tumor-induced axonogenesis. However, evidence supporting axonogenesis, particularly at early stages of the disease, came from studies on GEM models of PDAC, in which high densities of individual unbundled axons are observed in small regions of the pancreas containing fibrosis or pancreatic intraepithelial neoplasia (PanIN) lesions ^14^. These areas of hyperinnervation suggest that active axon growth may occur in a very localized manner. To address this, we employed LSFM to visualize the 3D distribution of TH-labeled sympathetic axonal endings within the pancreatic parenchyma. The main sympathetic nerve bundles described above defasciculated and splayed out into a meshwork of smaller bundles that entered each individual lobule of the pancreas. In control tissues, a dense network of fine sympathetic axonal fibers, which innervate the entire lobule, was observed (Fig. 2A–2C). Co-staining of blood vessels with MECA-32 or anti-PECAM antibodies indicated that TH^+^ fibers were aligned with the capillaries that supply blood to the acinar parenchyma, and on which they formed synapses as revealed by expression of the presynaptic marker synaptophysin 1 (Fig. 2D–2H). These results confirm previous observations of sympathetic innervation of the periacinar capillary bed ^19^. In pancreatic lobules of 6-week-old KIC mice, we observed “hot spots” of sympathetic hyperinnervation not present in control pancreata, while innervation of the rest of the tissue was similar to that of controls (Fig. 2I and 2J). These hot spots corresponded to dense networks of sympathetic fibers surrounding PanIN lesions, as revealed by autofluorescence signature analysis of the tissue (Fig. 2K and Supplementary Fig. S1). These findings indicate that changes in the sympathetic innervation of the pancreas occur early in the development of PDAC and may involve substantial growth and remodeling of individual nerve fiber terminals.

**Figure 2.**
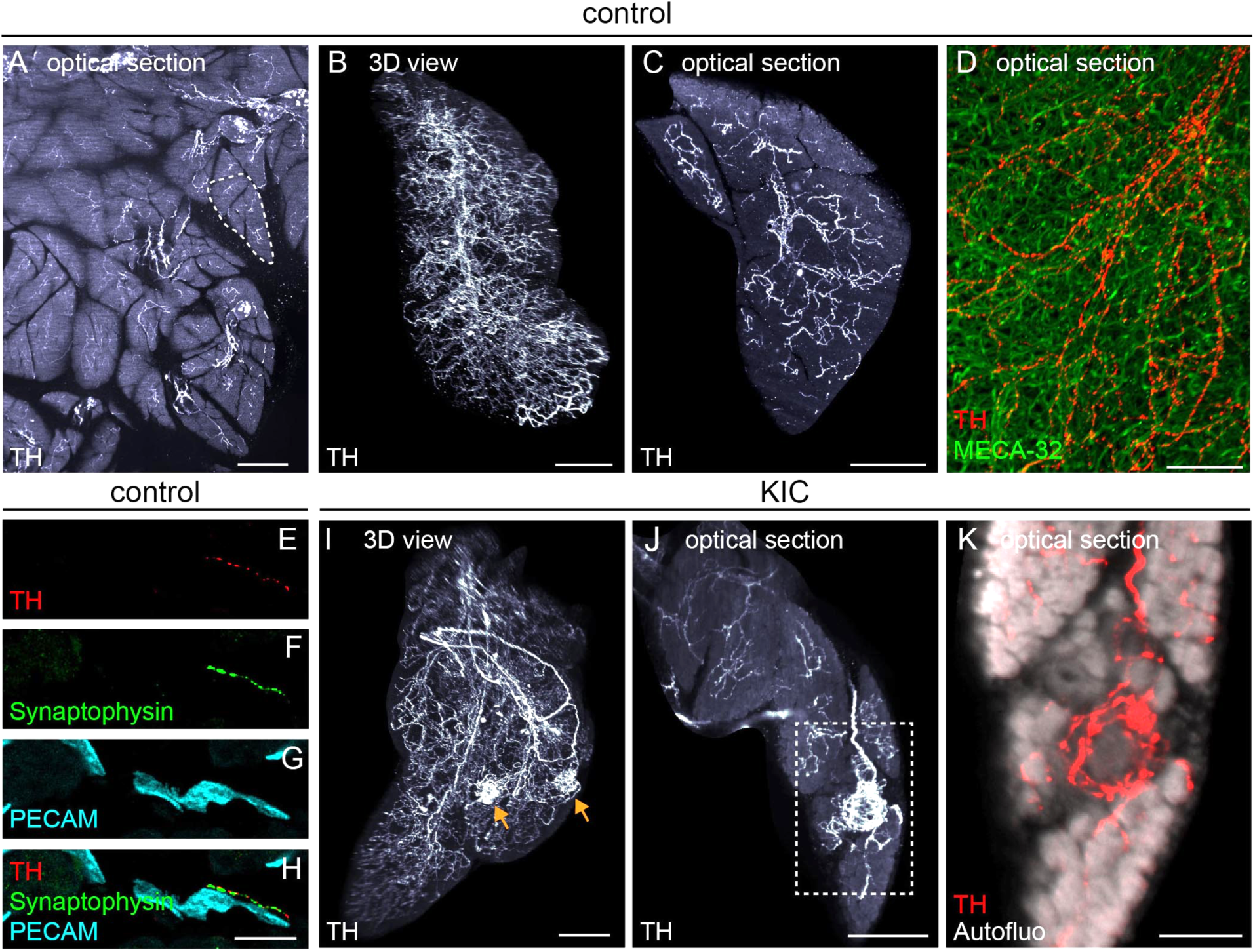
Hotspots of sympathetic innervation in KIC pancreas. **A**, Optical section through a cleared control pancreas immunostained with an anti-TH antibody. The dotted line delimits a pancreatic lobule. **B–C**, Representative 3D reconstruction (**B**) and optical section image (**C**) of a control pancreatic lobule immunostained for TH. **D**, Optical section through a pancreatic lobule doubled labeled with anti-TH and MECA-32 antibodies. **E–H**, Immunolabeling for TH (**E, H**), synaptophysin 1 (**F, H**), and PECAM (**G, H**) on a section through the acinar parenchyma of a control pancreas. **I–K**, 3D reconstruction (**I**) and optical section images (**J–K**) of a KIC pancreatic lobule showing “hotspots” of innervation by TH^+^ sympathetic axons (arrows). Tissue autofluorescence, imaged at an excitation wavelength of 488 nm, is shown in white **(K)**. Scale bars = 500 µm (**A**), 300 µm (**B**, **I**), 200 µm (**C**, **J**), 40 µm (**D**), 30 µm (**E–H**), and 100 µm (**K**).

### Sympathetic innervation of PDAC occurs through collateral axon sprouting

To thoroughly characterize how patterns of sympathetic nerve terminals change throughout successive stages of PDAC progression, we developed an image analysis pipeline that allows us to reconstruct axonal and vascular networks from 3D LSFM images. We analyzed a dozen parameters describing axon morphology and proximity interactions with blood vessels (Supplementary Fig. S2 and Supplementary Table S1). figs. 3A and 3B show original images and reconstructions of neuronal and vascular networks in acinar tissue of a control pancreas and a histologically normal (asymptomatic) region of a 6-week-old KIC pancreas. No major difference was observed, as the two tissues appeared similar in axonal and vascular density and the two systems were aligned next to each other, as evidenced by the reconstruction of their surfaces of contact. Moreover, the calculated axonal and vascular parameters were almost identical between controls and asymptomatic tissues with Z-scores close to 0 (Fig. 3G and Supplementary Table S2).

**Figure 3.**
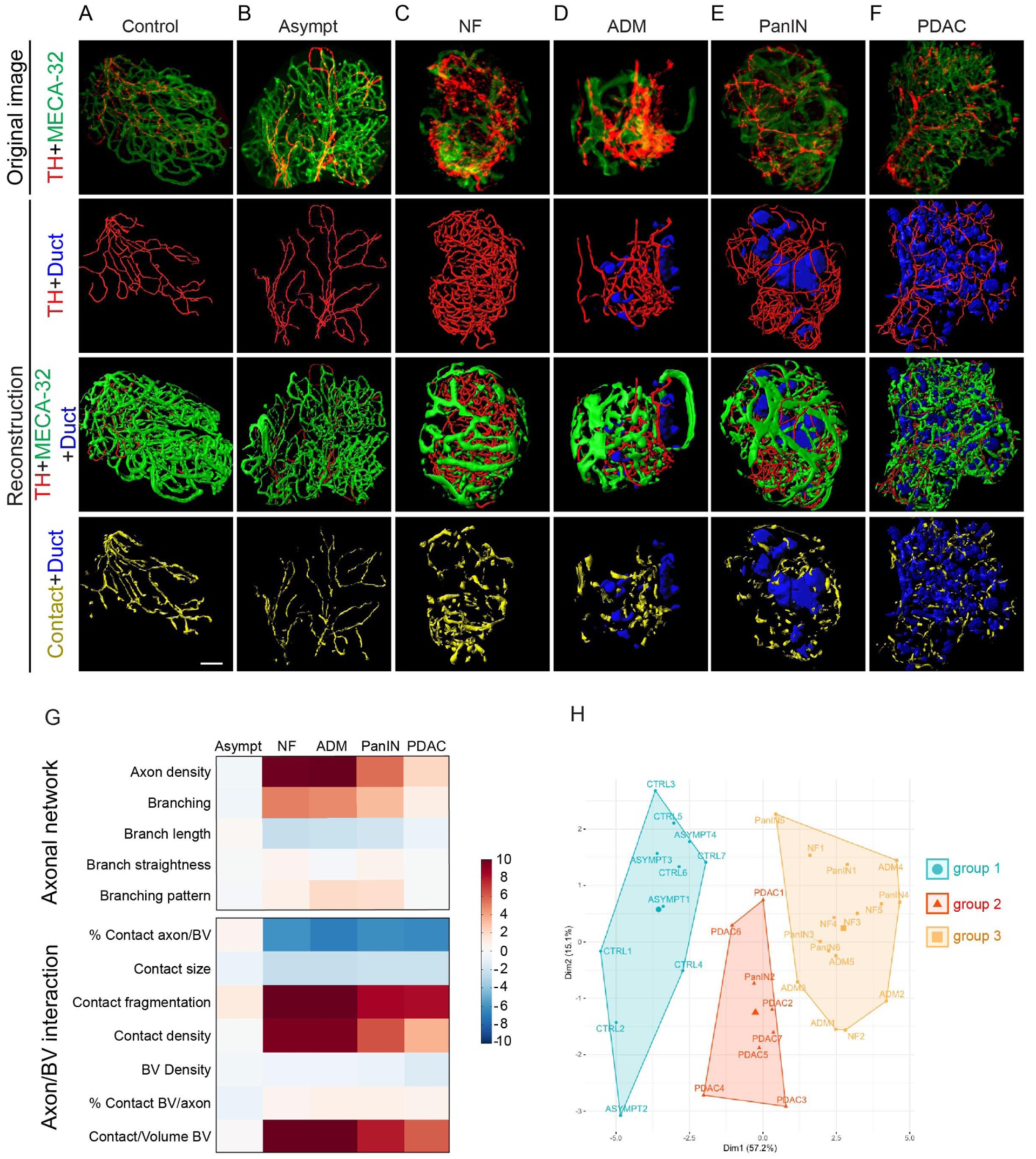
3D visualization and statistical analysis of sympathetic axon and blood vessel networks in KIC pancreas. **A–F**, Representative images of 8-week-old control (**A**) or KIC (**B–F**) pancreata immunostained with anti-TH and MECA-32 antibodies (first column). 3D reconstructions of sympathetic axons (red, second column), blood vessels (green, third column), and axon/vessel surface contacts (yellow, fourth column) in normal acinar tissue (**A**), asymptomatic acinar tissue (**B**), NF (**C**), ADM (**D**), PanIN (**E**), and a well-differentiated PDAC region (**F**). Cavities of the duct-like structures are represented in blue. **G**, Heatmap of the Z-scores calculated for each of the 12 variables describing the architecture of sympathetic axons and their relationship with blood vessels. Values from 4 asymptomatic (Asympt) acinar regions, 5 NF, 5 ADM, 6 PanIN, and 7 PDAC samples of 6-week-old KIC mice (n = 3) were compared with those of 7 normal acinar regions in age-matched control mice (n = 3). **H**, Factor map of the PCA performed on 34 tissue samples and 12 variables. Three cluster groups were identified corresponding to control and asymptomatic tissues (group 1, blue), non-invasive neoplastic pancreatic lesions (group 2, orange), and invasive tumor lesions (group 3, red). Scale bar = 50 µm.

On the other hand, sympathetic innervation was higher in pancreatic regions comprising nodular fibrosis (NF), acinar-to-ductal metaplasia (ADM), and PanIN lesions, which will be hereafter collectively referred to as “non-invasive pancreatic neoplastic lesions”, compared to control tissues (Fig. 3C–3E). Quantification confirmed increased densities of axons with more, but smaller, axon branches (Fig. 3G), indicating that hyperinnervation of these regions was due to localized sprouting of new collateral branches. Whereas sympathetic axons in healthy tissues were intimately associated with blood vessels, the contact surfaces between axons and vessels in non-invasive pancreatic neoplastic lesions appeared smaller (Fig. 3C–3E). Quantitative analysis of proximity contacts between remodeling axons and blood vessels showed an increased fragmentation level, with several smaller-sized contact zones (Fig. 3G). Overall, a lower percentage of sprouting axon surface area was in contact with blood vessels (% Contact axon/BV) compared with that of control conditions (Fig. 3G). This decrease could not be explained by decreased vascularization, as blood vessel density was not significantly altered between the examined regions—although blood vessels appeared structurally abnormal and enlarged in neoplastic lesions (Fig. 3C–3E). On the other hand, the percentage of vascular surface in contact with axons (% Contact BV/axon) remained unchanged across different regions, indicating that the reduction in axon/vessel contact size observed in the lesion regions may have been offset by the opposing increase in axonal sprouting and density.

Unbundled sympathetic axons were also detected in well-differentiated glandular regions of KIC tumors (Fig. 3F). These innervated PDAC regions were located on the periphery of tumor nodules, whereas undifferentiated regions in the tumor core were completely devoid of sympathetic nerve endings (data not shown). Well-differentiated PDAC regions had a slightly higher (but not statistically significant) sympathetic axonal density than that of control tissues and exhibited changes in axon-vessels contacts similar to those observed in non-invasive neoplastic pancreatic lesions (Fig. 3G).

Finally, using unsupervised hierarchical clustering and principal component analysis (PCA), we identified three different patterns of sympathetic innervation that distinguished the consecutive stages of PDAC progression from control and asymptomatic tissues (group 1) to non-invasive neoplastic lesions (group 2) and invasive PDAC (group 3) (Fig. 3H).

To further examine these findings, we performed similar experiments and analyses on LSL-Kras^G12D/+^; LSL-P53^R172H/+^; Pdx1-Cre (KPC) mice. KPC mice begin to develop PDAC at 8 weeks and have a median survival of about 5 months ^20^, which allowed us to study the innervation of tumors that developed over longer periods of time compared with that of KIC mice. In the pancreas of 14-week-old KPC mice, hyperinnervation of non-invasive neoplastic pancreatic lesions and changes in axon-vessel contacts similar to those reported in KIC mice were observed (Supplementary Fig. S3A–S3E and S3G). However, the axonal density measured in areas of well-differentiated PDAC was higher than that of the KIC tumors (Supplementary Fig. S3F–S3G and Supplementary Table S2). Therefore, in the KPC mouse model, PDAC clustered with the highly innervated non-invasive neoplastic pancreatic lesions in PCA (Supplementary Fig. S3H). Together, these data indicate that early and progressive sprouting and growth of sympathetic axons is a common characteristic of PDAC.

### Sprouting occurs from preexisting sympathetic axon fibers

A recent study on prostate cancer proposed that doublecortin (DCX)-expressing progenitor cells from the brain are transported via the bloodstream to tumors, where they contribute to the formation of new neurons and innervation of tumor tissues ^15^. We therefore assessed the presence of TH^+^ cell bodies in cleared samples and classical tissue sections of tumor-bearing KIC mice. No TH^+^ neuronal cell bodies were observed in normal acinar regions, consistent with the known organization of the sympathetic postganglionic innervation of the pancreas (data not shown). TH^+^ somata were detected in islets of Langerhans and corresponded to endocrine β-cells expressing insulin, as previously reported (Fig. 4A–4C) ^21,22^. Only rare scattered TH^+^ cell bodies were found in PDAC regions; these cells also expressed insulin and likely corresponded to the remains of islets destroyed by invasive tumor growth (Fig. 4D–4F). Meanwhile, many DCX^+^ cells were present in tumors but none were immunopositive for TH despite the presence of TH^+^ axonal fibers in the examined regions. In contrast, all DCX^+^ cells co-expressed the marker CD45 (Fig. 4G–4J), and thus most likely represent hematopoietic cells. Together, these results rule out a model of neurogenesis in PDAC and instead indicate that pancreatic tumors are innervated via the growth of pre-existing axon terminals.

**Figure 4.**
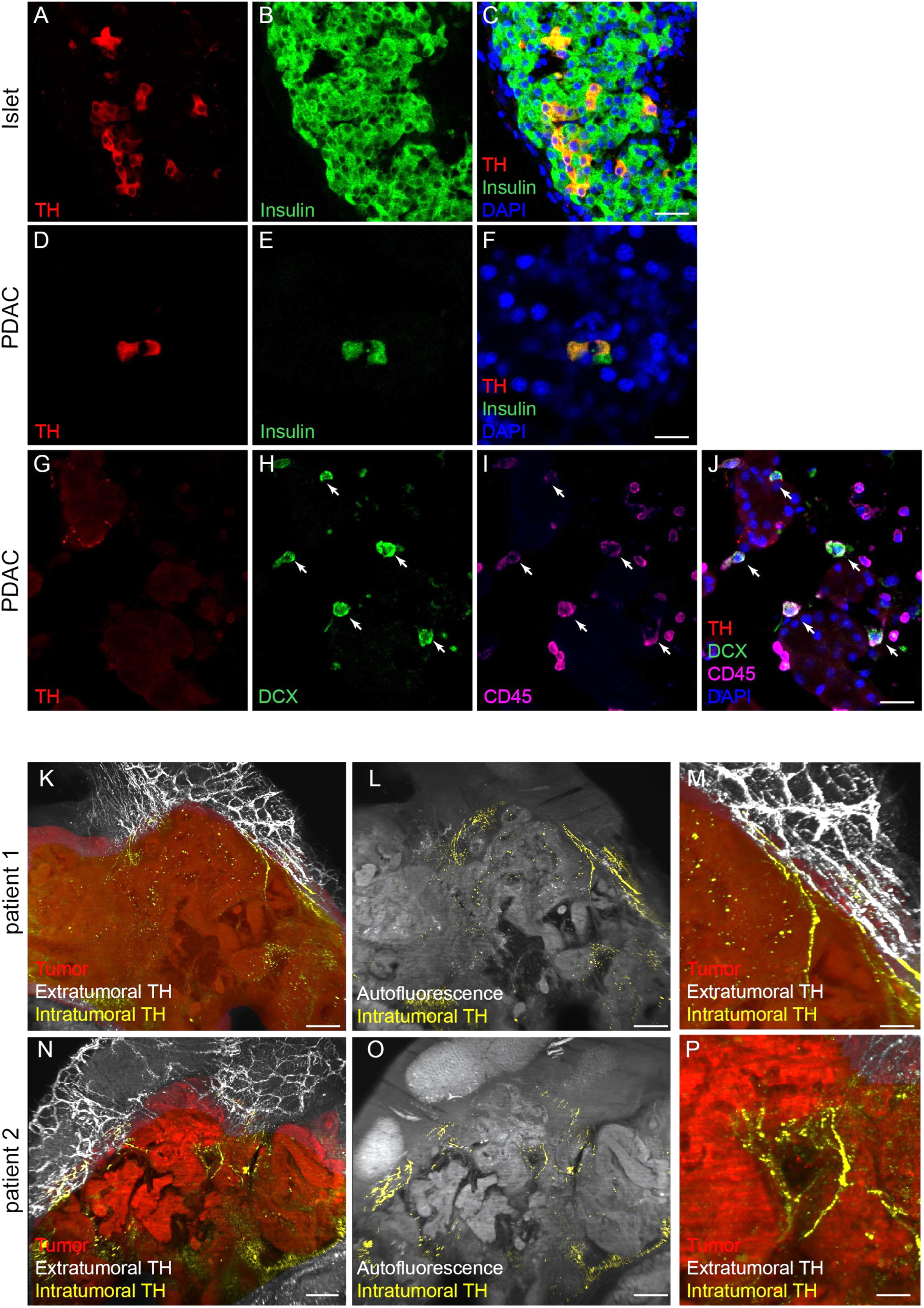
DCX expression in KIC tumors and sympathetic innervation of PDX models. **A–F**, Sections through the pancreas of 6-week-old KIC mice double labeled with anti-TH and anti-insulin antibodies. Images show an intact islet (**A–C**) and β-cells scattered inside the tumor (**D–F**). **G–J**, PDAC sections from 6-week-old KIC mice triple-labeled with anti-TH, anti-DCX and anti-CD45 antibodies. **K–P**, 3D view of PDX tumors derived from two different patients [patient 1 (**K–M)**, patient 2 (**N–P**)] and immunostained with anti-TH antibody. All panels are LSFM images of solvent-cleared tissue. 3D segmentation of PDX tumors is shown in red, intra-tumoral sympathetic fibers in yellow, and extra-tumoral fibers in white. Scale bars = 20 µm (**A–C** and **G–J**), 10 µm (**D–F**), 200 µm (**K**, **L**, **N**, **O**), and 50 µm (**M, P**).

To further test this notion, we employed orthotopic patient derived xenografts (PDXs) to assess the ability of graft tumors to recruit host sympathetic axon terminals. We used two defined subtypes of PDX tumors pre-characterized as “classical” and “basal-like” in a previous multiomic analysis ^23^. LSFM imaging of TH-labeled axons in undifferentiated basal-like PDXs revealed a lack of sympathetic innervation (n = 4 PDX tumors derived from two human patients), reflecting the absence of fibers in the undifferentiated regions of KIC tumors (data not shown). In contrast, TH-labeled fibers were detected in classical PDXs (n = 4 PDX tumors derived from two human patients). Virtual dissection and 3D reconstruction of the projection networks revealed that the observed intra-tumoral sympathetic axons were distal extensions of fibers that innervated the adjacent regions of the murine tissue (Fig. 4K–4P). A clear demarcation between PDX and mouse pancreatic tissue was always visible, which excluded a simple engulfment of mouse fibers. These data confirm that both murine and PDX tumors are capable of stimulating growth and recruiting pre-existing sympathetic axons from surrounding tissues.

### Sympathectomy accelerates tumor growth and metastasis and decreases survival

The neuroplastic changes described are likely to play a significant role in PDAC progression. To test this, we used the neurotoxin 6-hydroxydopamine (6-OHDA) to induce peripheral chemical sympathectomy in KIC mice (Fig. 5A). Mice received two intraperitoneal (i.p.) injections of 6-OHDA or ascorbic acid (AA) vehicle solution between 3 and 4 weeks of age, i.e., after the sympathetic-dependent maturation of endocrine pancreatic functions and before premalignant lesions develop ^17,21^. 6-OHDA treatment selectively eliminated 83.9 % of TH^+^ axons residing in acinar tissue, without affecting vesicular acetylcholine transporter (VAChT)-positive parasympathetic fibers or blood vessel density (Supplementary Fig. S4A-S4G). We found that 6-OHDA treatment significantly reduced KIC mouse survival compared with vehicle-treated mice (median survival, KIC^AA^: 9.1 weeks, KIC^6-OHDA^: 7.8 weeks; Fig. 5B). Chemical sympathectomy had no effect on the survival of non-tumor bearing mice during the same period of time (Fig. 5B). Notably, KIC mice rarely developed macrometastatic disease ^17^, whereas autopsy at death revealed high percentages of 6-OHDA-treated mice with hepatic metastasis (KIC^AA^: 16.7 %, KIC^6-OHDA^: 77.8 %) or peritoneal carcinomatosis (KIC^AA^: 8.3 %, KIC^6-OHDA^: 77.8 %) (Fig. 5C–5J).

**Figure 5.**
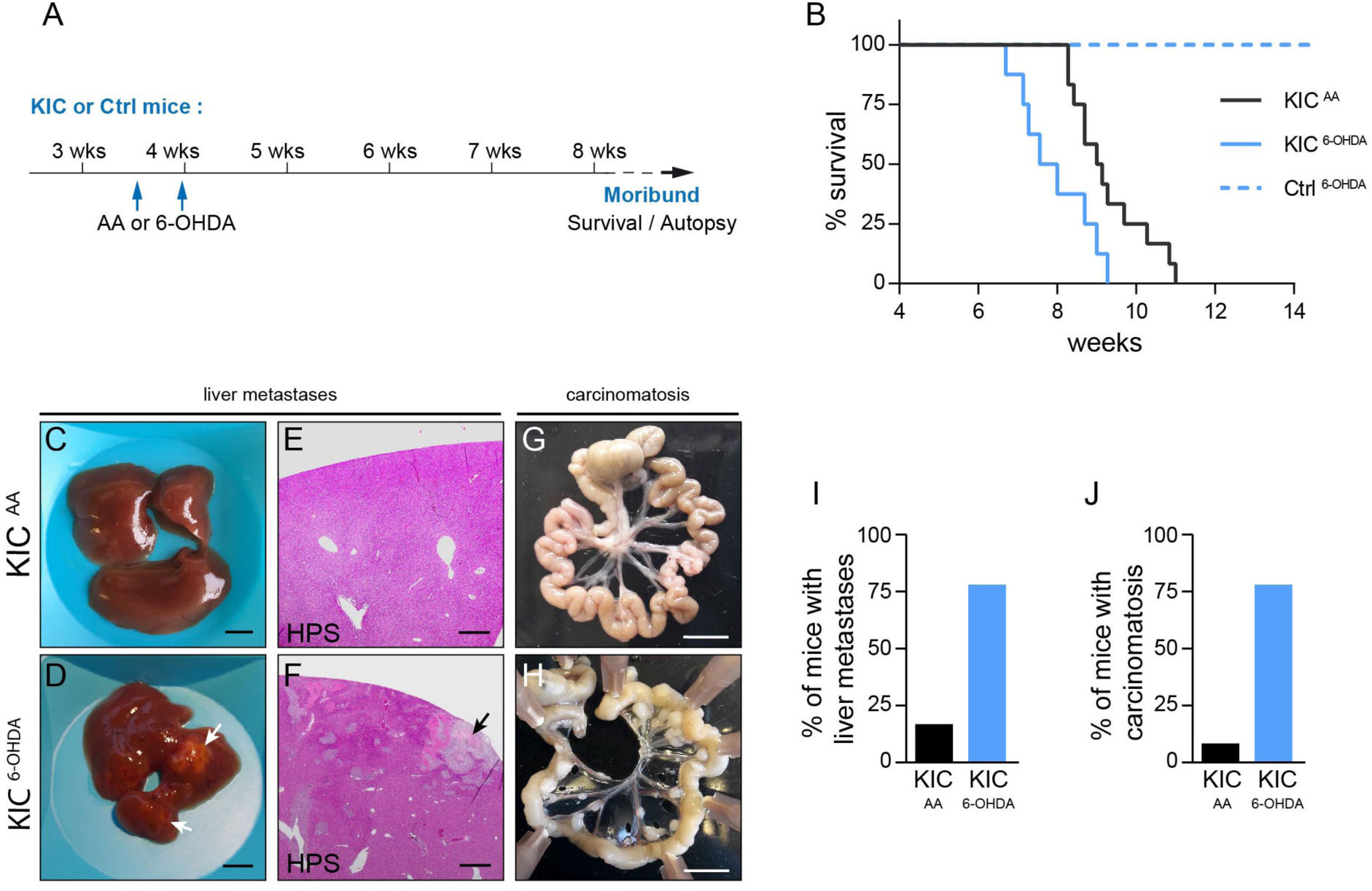
Reduced survival and increased metastatic spread in sympathectomized KIC mice. **A**, Outline of the experiment. **B**, Kaplan-Meier curve comparing overall survival of control mice treated with 6-OHDA (Ctrl6-OHDA, n = 10) and KIC mice treated with AA (KIC^AA^, n = 12) or 6-OHDA (KIC^6-OHDA^, n = 8). For KIC^AA^ vs. KIC^6-OHDA^: log-rank = 0.0132 and hazard ratio B/A = 2.711 (A = KIC^AA^ and B = KIC^6-OHDA^). **C–H**, Representative pictures of livers (**C, D**), HPS-stained liver sections (**E, F**), and intestine (**G, H**) of AA- or 6-OHDA-treated KIC mice collected at moribund stages. The white arrows point to macrometastases. **I–J**, Graph showing the percentage of AA- and 6-OHDA-treated KIC mice with liver metastasis (**I**) and carcinomatosis in the intestine mesentery (**J**) at death (KIC^AA^, n = 9; KIC^6-OHDA^, n = 6). Scale bars = 5 mm (**C, D** and **G, H**), and 500 µm (**E** and **F**).

To further assess these results, we performed surgical sympathectomy by severing the nerves entering the pancreas, which contained a mix of sympathetic and sensory fibers (Supplementary Fig. S4C-S4J). Compared with sham-operated animals, KIC mice that underwent pancreatic denervation showed decreased survival (median survival, KIC^Sham^: 9.0 weeks, KIC^SympX^: 8.1 weeks, Supplementary Fig. S4K-S4L) and a higher proportion of liver metastases (KIC^Sham^: 33.3 %, KIC^SympX^: 66.7 %) and carcinomatosis (KIC^Sham^: 11.1 %, KIC^SympX^: 33.3 %) (Supplementary Fig. S4M–S4R).

Next, to precisely analyze the effect of the sympathetic nervous system on primary tumor growth, we performed chemical sympathectomy of KIC mice with 6-OHDA at ^3–4^ weeks and comprehensively analyzed tumors from animals at 6.5 weeks when the tumors became palpable (Fig. 6A). Total pancreatic weight increased in sympathectomized KIC mice compared with vehicle-treated KIC animals (Fig. 6B–6D). Pathological evaluation of the denervated pancreas revealed that the area occupied by the tumor increased by 30 % at the expense of histologically normal tissue (Fig. 6E–6G). These larger tumors were associated with increased cell proliferation (Fig. 6H, 6I, and 6P), increased fibrosis (Fig. 6J, 6K, and 6Q), reduced vascular density (Fig. 6L, 6M, and 6R), and increased hypoxia (Fig. 6N, 6O, and 6S). All these changes are indicative of a more advanced or aggressive malignancy.

**Figure 6.**
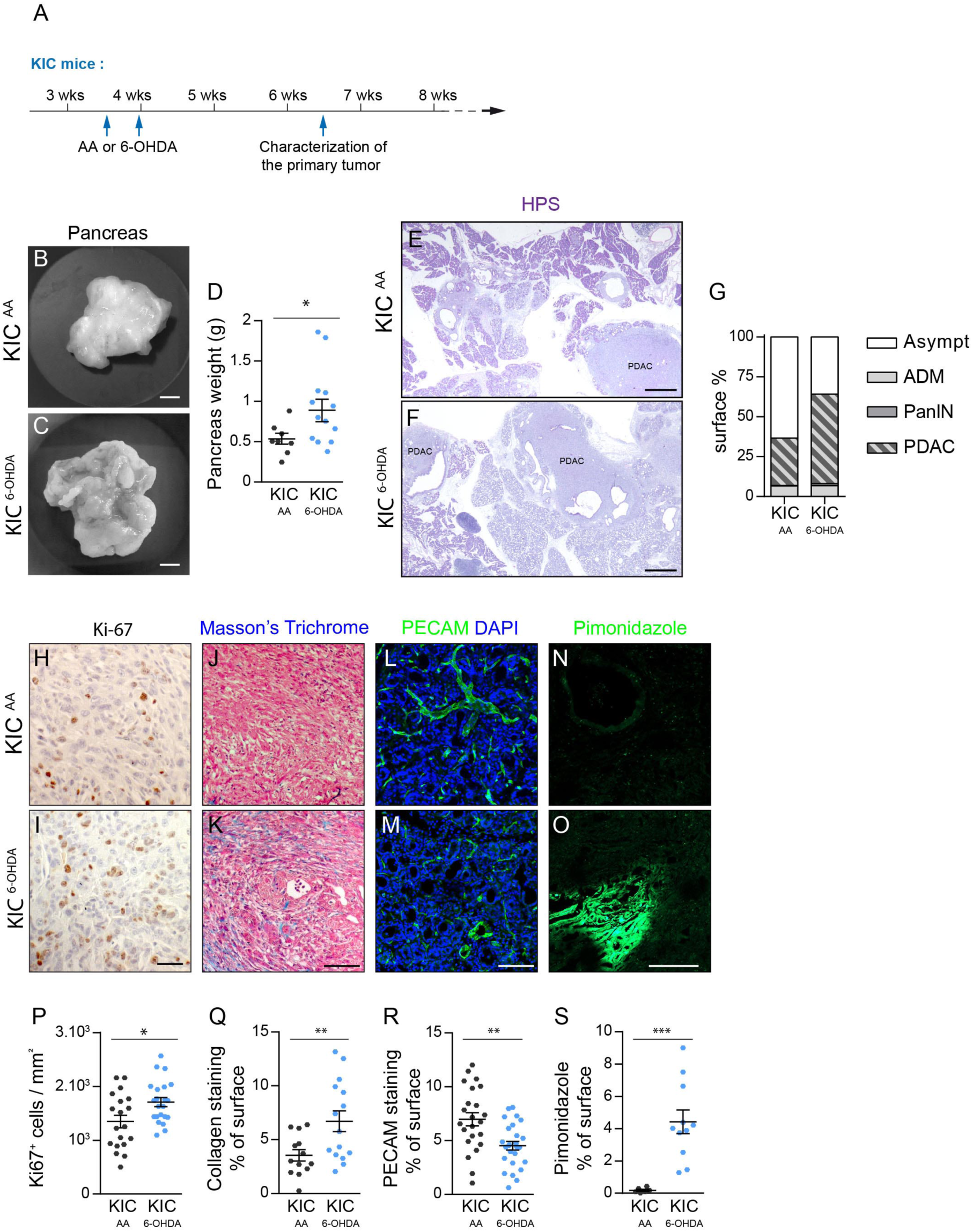
Accelerated tumor growth in sympathectomized KIC mice. **A**, Outline of the experiment. **B**, **C**, Representative pictures of tumoral pancreata from 6.5-week-old KIC mice treated with AA (**B**) or 6-OHDA (**C**). **D**, Scattered dot plot of pancreatic weight from 7-week-old KIC mice treated with AA (KIC^AA^, n = 9) or 6-OHDA (KIC^6-OHDA^, n = 13). p = 0.0348 (unpaired *t*-test with Welch’s correction). **E, F**, Bright field images of HPS-stained pancreatic sections from 6.5-week-old KIC mice treated with AA or 6-OHDA. **G**, Stacked bar graph showing the percentage of surface covered by asymptomatic tissue (Asympt), ADM, PanIN, and PDAC in pancreatic sections from 6.5-week-old KIC mice treated with AA (n = 3 mice, 12 sections) or 6-OHDA (n = 3 mice, 12 sections). Asympt, p = 0.0447 (Mann– Whitney test); ADM, p = 0.9907 (unpaired *t*-test); PanIN, p = 0.1088 (Mann–Whitney test); PDAC, p = 0.0177 (unpaired *t*-test). **H–O**, Representative images of immunohistochemistry for Ki-67 (**H, I**), Masson’s trichrome staining (**J, K**), immunofluorescence for PECAM (**L, M**), and pimonidazole staining (**N, O**) in PDAC sections from 6.5-week-old KIC mice treated with AA or 6-OHDA. **P–S**, Scattered dot plots of the densities of Ki-67+ cells (**P**), collagen (**Q**), PECAM+ vessels (**R**), and pimonidazole+ hypoxic areas (**S**) in PDAC sections of 6.5-week-old KIC mice treated with AA or 6-OHDA. **P**, p = 0.0145, unpaired *t*-test with Welch’s correction (KIC^AA^, n = 3 mice, 19 images; KIC^6-OHDA^, n = 3 mice, 22 images). **Q**, p = 0.0085, unpaired *t*-test with Welch’s correction (KIC^AA^, n = 3 mice, 13 images; KIC^6-OHDA^, n = 3 mice, 15 images). **R**, p = 0.0014, unpaired *t*-test (KIC^AA^, n = 6 mice, 23 images; KIC^6-OHDA^, n = 5 mice, 26 images). **S**, p = 0.0002, Mann–Whitney test (KIC^AA^, n = 1 mouse, 6 images; KIC^6-OHDA^, n = 2 mice, 11 images). Scale bars = 5 mm (**B**, **C**), 500 µm (**E**, **F**), 50 µm (**H**, **I**), 100 µm (**J**, **K**), and 200 µm (**L–O**). Error bars represent median ± SEM

The promotive effect induced by sympathetic denervation on tumor growth was further tested in a syngeneic orthotopic mouse model of PDAC. For this, luciferase-expressing epithelial cancer cells (PK4A-Luc cells) isolated from a KIC tumor ^24^ were injected into the pancreas of 6-OHDA-lesioned or vehicle-treated mice (Supplementary Fig. S4K). Longitudinal monitoring of tumor development via bioluminescence and statistical analysis revealed that initial tumor growth rate was higher in sympathectomized mice compared with controls (Supplementary Fig. S4S and S4T). Altogether, these results demonstrate that the sympathetic nervous system exerts a protective function during development and progression of pancreatic cancer.

### Sympathectomy increased intra-tumoral macrophages

To elucidate the mechanism underlying the observed effects of sympathetic nerve ablation, we first examined immune infiltration of tumors. We focused in particular on macrophages whose function can be modulated by the sympathetic nervous system in several organs and disease contexts ^25^. To label macrophages in pancreatic sections of chemically sympathectomized or control KIC mice, we used the macrophage-specific markers F4/80 and CD163, the latter being frequently observed in aggressive tumors ^26^. All cells expressing the surface receptor CD163 were also F4/80^+^ and accounted for nearly half of the macrophages found in asymptomatic pancreas regions (Supplementary Fig. S5A–S5D). In these same regions, administration of 6-OHDA did not alter the density of CD163^+^ or F4/80^+^ macrophages (Supplementary Fig. S5B– S5D). In contrast, a significant increase in CD163^+^ macrophage number was observed in non-invasive pancreatic lesions (ADM and PanIN) of 6-OHDA-treated pancreata, while the total number of F4/80 macrophages remained constant (Supplementary Fig. S5E–S5L). Moreover, F4/80^+^ macrophages were present in higher numbers in PDAC of sympathectomized mice compared with control tumors (Fig. 7A, 7B, and 7D). Sympathectomy was also associated with an important change in the distribution of CD163^+^ macrophages that appeared to infiltrate tumors (Fig. 7A, 7B), whereas this macrophage population was largely found in the peritumoral region of control tumors (not shown). These observations were supported by the increase in intratumoral CD163 expression levels and CD163/F4/80 ratio in 6-OHDA-treated PDAC compared with controls (Fig. 7C–7E). Together, these findings indicate that sympathectomy induces a high proportion of CD163^+^ macrophages in both preneoplastic and tumor tissues.

**Figure 7.**
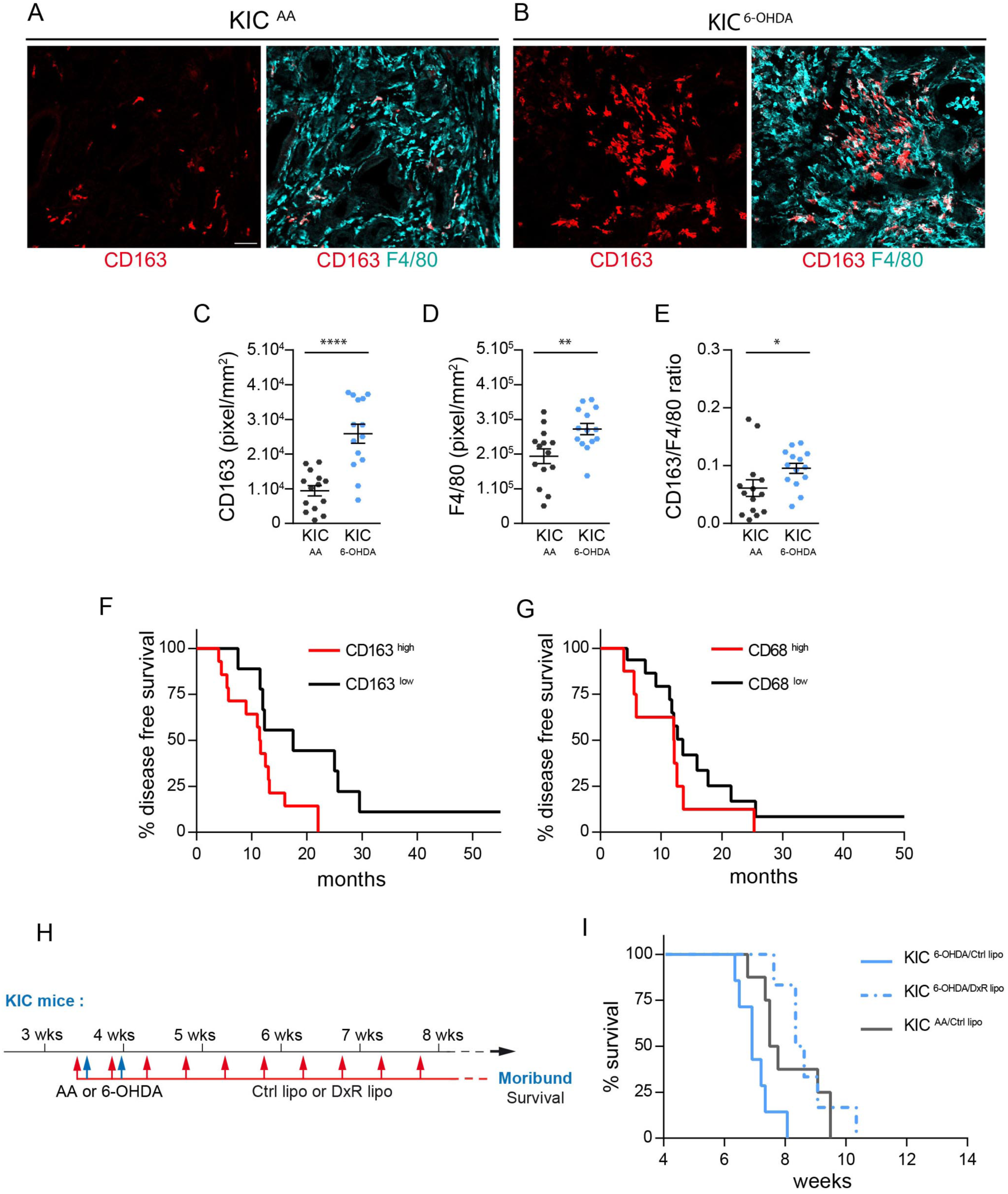
CD163^+^ macrophages mediate the effect of sympathectomy. **A–B**, Representative images of F4/80^+^ and CD163^+^ macrophages in PDAC of 6.5-week-old KIC mice treated with AA (**A**) or 6-OHDA (**B**). The boxed areas are enlarged to show CD163**+** macrophage infiltration in tumors of sympathectomized mice. **C–E**, Dot plots depicting the densities of CD163^+^ macrophages (**C**; p < 0.0001, unpaired *t*-test with Welch’s correction), F4/80+ macrophages (**D**; p = 0.0073, unpaired *t*-test), and the CD163/F4/80 ratio (**E**; p = 0.0141, Mann–Whitney test) in PDAC of 6.5-week-old KIC mice treated with AA (KIC^AA^, n = 4 mice, 14 images) or 6-OHDA (KIC^6-OHDA^, n = 3 mice, 14 images). Error bars represent median ± SEM. **F**, Kaplan–Meier curves comparing disease-free survival of human patients with PDAC and high intratumoral CD163 (n = 14) or low CD163 (n = 9) expression levels. Log-rank = 0.0231; hazard ratio A/B = 3.428 and B/A = 2.445 (A = low CD163 and B = high CD163). **G**, Kaplan–Meier curve comparing overall survival of human patients with PDAC and high intratumoral CD68 (n = 8) or low CD68 (n = 17) expression levels. Log-rank = 0.1838; hazard ratio A/B = 0.5552 and B/A = 1.801 (A = low CD68 and B = high CD68). **H**, Outline of the experiment quantified in (**I**). **I**, Kaplan–Meier curve comparing overall survival of AA-treated KIC mice injected with Ctrl lipo (KIC^AA/Ctrl lipo^, n = 8) and 6-OHDA-treated KIC mice injected with Ctrl lipo (KIC^6-OHDA/Ctrl lipo^, n = 7) or DxR lipo (KIC^6-OHDA/DxR lipo^, n = 6). KIC^AA/Ctrl lipo^ vs. KIC^6-OHDA/Ctrl lipo^, log-rank = 0.0244 and hazard ratio B/A = 2.778 (A = KIC^AA/Ctrl lipo^ and B = KIC^6-OHDA/Ctrl lipo^). KIC^6-OHDA/Ctrl lipo^ vs. KIC^6-OHDA/DxR lipo^, log-rank = 0.0011 and hazard ratio A/B = 4.173 (A = KIC^6-OHDA/Ctrl lipo^ and B = KIC^6-OHDA/DxR lipo^). Scale bars = 500 µm (**A**, **B**) and 150 µm (insets).

### Depletion of CD163^+^ macrophages rescued the effect of sympathectomy

Given the association between high macrophage infiltration and poor outcome in most cancers, we reasoned that the changes observed in the number and/or type of tumor-associated macrophages (TAMs) may contribute to the more severe phenotypes in sympathectomized KIC mice. In an analysis of macrophage populations in PDAC specimens from 34 patients treated at the North Hospital of Marseille (France), we found that high intra-tumoral expression of CD163, but not of the pan-macrophage marker CD68, was associated with shorter disease-free survival (CD163^low^: 17.5 months, CD163^high^: 11.5 months; Fig. 7F and 7G), confirming a previous report ^27^. Based on these findings, we therefore decided to test the role of CD163^+^ macrophages in the survival of sympathectomized KIC mice. To this end, we used polyethylene glycol-coated liposomes vectorized with CD163 antibody to selectively deliver the chemotoxic agent doxorubicin (DxR lipo) to CD163^+^ macrophages ^28^. Liposome administration via the retro-orbital route in KIC mice led to targeted depletion of 68.5 % of the CD163^+^ TAMs in PDAC, whereas control liposomes non-loaded with doxorubicin (Ctrl lipo) exhibited no effects, and increased mouse survival (Supplementary Fig. S5M–S5R). Sympathectomized KIC mice were treated with either DxR lipo or Ctrl lipo three times per week until death due to tumor burden (Fig. 7H). Ablation of CD163^+^ macrophages improved survival of sympathectomized mice (median survival, KIC^6-OHDA/Ctrl lipo^: 6.8 weeks, KIC^6-OHDA/DxR lipo^: 8.4 weeks), which was restored to the same level as non-denervated KIC mice (Fig. 7I). Taken together, these findings support a role for CD163^+^ macrophages as mediators of the pro-tumoral effect of sympathectomy in PDAC.

## Discussion

A typical feature of pancreatic carcinoma is the presence of intra-tumoral nerves, often hypertrophic and in greater numbers than in normal pancreatic tissue ^29^. Pancreatic cancer is therefore thought to exert neurotrophic properties that induce the formation and growth of new nerves into the tumor. Contrary to this model, the present study showed that at least some of the sympathetic nerves found inside PDAC tumors are pre-existing internal pancreatic nerves. These nerves have been engulfed by the tumor and their presence is therefore not the result of an active process of nerve growth and plasticity. 3D imaging highlighted the heterogenous distribution of sympathetic nerves within the pancreas. Thus, nerve density in tumor samples may vary considerably depending on tumor location. Moreover, in experimental PDAC tumors, increased sympathetic nerve density was reported after treatments causing a concomitant increase in tumor development ^8^. However, compared with small tumors, larger tumors that have invaded most of the organ are more likely to contain embedded pre-existing nerves. Thus, tumor size and location are parameters that may account for differences in intra-tumoral nerve density, without the growth of new nerves being involved. This is a critical point that should be considered in studies evaluating the prognostic value of intra-tumoral nerve density in cancer.

While sympathetic nerves do not appear to undergo active changes, we found a significant increase in terminal axon density in non-invasive neoplastic lesions and, to a lesser extent, in well-differentiated regions of PDAC tumors. It is unlikely that these axons emerged from new intra-pancreatic neurons generated by brain-derived precursors as proposed ^15^, as we found no TH^+^ neuronal cell bodies in the pancreas and tumors. We did, however, detect DCX-expressing cells in pancreatic tumors. In addition to being a neurogenesis marker, the microtubule-associated protein DCX is also expressed outside neurogenic niches by hematopoietic cells, including CD8^+^ T cells ^30^. We therefore propose that the DCX^+^/CD45^+^ cells observed in PDAC correspond to infiltrating immune cells and do not reflect intra-tumoral neurogenesis.

Sympathetic innervation of pancreatic lesions would therefore result from sprouting of new axon collaterals from axon terminals that normally innervate blood capillaries located close to the lesioned areas. This is supported by our data showing an increased number of short axonal branches in both non-invasive neoplastic and invasive pancreatic lesions. Sprouting responses from sympathetic axons has been documented in several chronic inflammatory diseases of the skin, joints, and mucosa ^31–33^. PDAC is a typical example of an inflammation-linked cancer. The persistent chronic micro-inflammation induced by oncogenic *Kras* that drives the initiation of precancerous lesions ^34^ may be at the origin of abnormal axonal sprouting and sympathetic hyperinnervation observed in both KIC and KPC pancreas. The mechanisms linking inflammation to sympathetic neural remodeling remain to be fully elucidated, though a role for NGF has been proposed ^35^. Sprouting may also be regulated by axon guidance molecules. Indeed, genomic alterations in axon guidance genes (including Slit/Robo and Semaphorin pathways) in human patients with PDAC as well as increased expression in mouse models of early neoplastic transformation and invasive PDAC have been identified ^36^. Some axon guidance molecules can regulate adult sympathetic axon growth in cell culture systems, but their involvement in innervation of pancreatic lesions is yet to be demonstrated in vivo.

Our results revealed an intra-tumoral heterogeneity in the distribution of axonal sprouts, which appears correlated with the degree of histological tissue differentiation. Indeed, sympathetic axon terminals were restricted in well-differentiated glandular areas, often located at the periphery of tumor foci, while undifferentiated regions of tumors with dense desmoplastic stroma were completely devoid of infiltrated nerve endings, suggesting that these regions may be non-permissive and/or too compact for significant axon growth. Interestingly, previous PDAC classification studies identified at least two main clinically relevant subtypes, which also differ in terms of histological differentiation grades; classical tumors are more often well-differentiated and basal tumors are more often poorly differentiated ^37^. The present finding that only “classical” PDX tumors were able to attract host sympathetic axon growth suggests that different subtypes of human PDAC may have distinct neural environments that contribute to their different clinical outcomes.

The present study revealed a protective role of the sympathetic nervous system against PDAC. This is reminiscent of the beneficial effect the sympathetic nervous system exerts on pancreatic tumor development in mice exposed to eustress ^10^. In the same study, chemical sympathectomy also showed a tendency to promote transplanted pancreatic tumor growth in mice reared under standard environments. However, the reduced survival of sympathectomized KIC mice contrasts with a recent study in which sympathectomy, alone or combined with gemcitabine, was associated with longer overall survival in the transgenic KPC model ^8^. This inconsistency may be explained by the time point at which denervation was performed—before disease onset in the present study and after tumor establishment in the other case. The temporality of denervation may have a strong influence on tumorigenesis, as has been reported for prostate cancer, where experimental sympathectomy has an effect only when performed before invasive disease onset ^3^. Pancreatic sympathectomy before disease onset may prevent the early remodeling of sympathetic endings that leads to hyperinnervation of ADM and PanIN lesions. The resulting worsening of KIC mouse phenotypes suggests that the sympathetic nervous system may slow the progression of precursor lesions toward the invasive form of the disease.

On the other hand, sympathectomy during invasive PDAC may bypass this early function, instead revealing a later tumor growth-promoting function of the sympathetic nervous system. Such a function, however, is not supported by our bioluminescent imaging of grafted tumors, which instead revealed an inhibitory function of the sympathetic nervous system on tumor growth. As the late denervation experiments were performed by surgical resection of nerves around the celiac-superior mesenteric ganglion complex ^8^, the observed effects may more probably result from elimination of the sensory component of these mixed sympathetic-sensory nerves. Indeed, previous studies reported that selective depletion or inactivation of sensory nerve fibers reduced tumor growth and prolongated overall survival in KPC mice ^38,39^.

While early sympathetic denervation inhibits pancreatic tumorigenesis, an opposite accelerating effect on tumor development has been reported in several other cancer types, including prostate, breast, and liver cancers ^3,40,41^. A determining factor could be the type of sympathetic neurons involved. A recent single-cell RNA-seq study of sympathetic neurons from the thoracic ganglion chain revealed the existence of five subtypes of TH-expressing neurons ^42^. These data indicate a greater degree of diversity and functional specialization of sympathetic neurons than previously thought. Whether this could explain their different activities during tumorigenesis awaits further investigation.

Alternatively, divergent tumoral responses to early sympathetic ablation may depend on the specific type of cancer studied. In prostate cancer, increased sympathetic axon density is accompanied by increased axon/blood vessel interactions, while adrenergic signaling to endothelial cells promotes angiogenesis and thus cancer growth ^43^. In pancreatic cancer, the sprouting of sympathetic axons is also likely to stimulate angiogenesis through increased local paracrine signaling to endothelial cells. In agreement, we found that sympathectomized PDAC tumors displayed reduced intra-tumor vascularization. While the primary angiogenic function of the sympathetic nervous system may be a common feature in cancer, decreased tumor vascularity after denervation, however, may not have the same impact in PDAC as in other cancers. Indeed, PDAC is typically a hypovascular tumor—due to its desmoplastic stroma—that is well adapted to hypoxic conditions ^44^. Reduced vascularity, which is a favorable factor in some cancers, is on the contrary associated with poorer survival in PDAC ^45^. Thus, PDAC response to sympathectomy may be shaped by its particular microenvironment.

Lastly, we identified a crucial function of the sympathetic nervous system in controlling the tumor immune environment. In KIC mice, pancreatic sympathectomy induced an increase in CD163^+^ macrophage density, which begins during the earliest stages of cancer progression, and later infiltration into PDAC tumors. Importantly, this “reprogramming” of macrophages mediated the pro-tumor response to pancreatic sympathectomy, as selective depletion of the CD163^+^ TAM population—which represents only a small proportion (around 10%) of all TAMs—was sufficient to rescue overall survival of sympathectomized KIC mice. The ability of CD163^+^ TAMs to promote tumor progression has recently been demonstrated in mouse models of melanoma, where the CD163^+^ TAM subset plays a key role in suppressing both myeloid and T cell-mediated anti-tumor immunity ^28^. Thus, how can sympathetic afferents of the pancreas help prevent macrophages from adopting a tumor-promoting phenotype? Sympathetic axon sprouting in PDAC and its precursor lesions may result in a local increase in the release of norepinephrine and its co-transmitters, all of which can have a direct influence on immune cells. The effect of norepinephrine has been well characterized and shows a modulating activity on macrophages (toward pro- or anti-inflammatory phenotypes) that varies considerably with context ^46^. Thus, it is likely that sympathetic nerve activity suppresses CD163^+^ macrophages while activating other subsets of macrophages or immune cells. Nevertheless, an indirect effect on macrophages, which can adopt pro-tumor phenotypes in response to sympathectomy-induced tumor hypoxia ^47^, cannot be excluded. Future comprehensive studies investigating the immune landscape of control and denervated PDAC will determine how sympathetic afferents of the pancreas contribute to a protective immune environment.

In conclusion, we demonstrated that sympathetic innervation of non-invasive and invasive PDAC lesions occurs independently of neurogenesis via the sprouting of sympathetic axonal endings that normally supply the exocrine pancreatic capillaries. Moreover, we showed that sympathetic axons slow pancreatic tumor progression through local suppression of CD163^+^ macrophage subsets at lesion sites. Recent studies reported a similar inhibitory role of the parasympathetic nervous system in PDAC, which is exerted in part by the suppression of myeloid cells ^6,7^. Tumor formation and progression have been widely regarded as dependent on autonomic nerve function, stimulating interest in clinical investigation of inhibitors of adrenergic and/or muscarinic cholinergic signaling in the treatment of certain cancers. However, our findings indicate a higher level of functional diversity of autonomic nerves in cancer than previously thought and suggest that blocking autonomic nerve function is not a therapeutic option for PDAC. The challenge is to exploit these new insights into autonomic nervous system regulation of PDAC immune infiltrate and translate it into clinical strategies.

## Methods

### Mouse strains

All animal procedures were conducted in accordance with the guidelines of the French Ministry of Agriculture (approval number F1305521) and approved by the local ethics committee (CE14 approval number APAFIS#1325-2016120211301815v1, APAFIS#17278-2018102514126851V3, and APAFIS#17026-2018092610362806 v6). Wild-type mice had a C57BL/6 background or a mixed FVB/C57BL/6 background (Janvier Labs, Le Genest-Saint-Isle, France), while NU/NU nude mice were obtained from Charles River Laboratories (Wilmington, MA). KIC (LSL-Kras^G12D/+^; Ink4a/Arf^lox/lox^; Pdx1-Cre) and KPC (LSL-Kras^G12D/+^; LSL-P53^R172H/+^; Pdx1-Cre) mutant mice were obtained by intercrossing LSL-Kras^G12D/+^ ^48^, Pdx1-Cre ^49^, and Ink4a/Arf^lox/lox^ ^17^ or LSL-P53^R172H/+^ ^20^ mice.

### Whole-mount immunostaining and clearing procedure

Intact pancreata were immunostained and cleared following the iDISCO^+^ protocol ^50^. The antibodies used are listed in Supplementary Table S3. In some experiments, nuclear counterstaining was performed by incubation in TO-PRO™-3 Iodide (642/661; cat. #T3605; Thermo Fisher Scientific, Waltham, MA) diluted in DMSO (1:1000) for 90 min at room temperature after incubation with secondary antibodies.

### LSFM and image processing

3D imaging was performed with a light sheet fluorescence microscope (Ultramicroscope II; Miltenyi Biotec, Bergisch Gladbach, Germany) using Imspector Pro software (Miltenyi Biotec). 3D volume images were generated using Imaris x64 software (version 9.2.1 and 9.3.0; Bitplane, Zurich, Switzerland). Stack images were first converted to Imaris files (.ims) using Imaris FileConverter. 3D reconstruction of samples was then performed using “3D view” in Imaris. The samples could be optically sliced in any angle using the “Orthoslicer” or “Obliqueslicer” tools. 3D pictures and movies were generated using the “Snapshot” and “Animation” tools. Finally, images were cropped and, if required, their brightness adjusted evenly using Photoshop CS6 (Adobe, San Jose, CA). The pipeline for quantitative image analysis is recapitulated in Supplementary Fig. S2 and Supplementary Table S1.

### Denervation experiments

For chemical sympathectomy, 6-hydoxydopamine (6-OHDA; cat. #H4381; Sigma-Aldrich, St. Louis, MO) was dissolved in vehicle solution (0.1 % ascorbic acid [cat. #A4403; Sigma-Aldrich]) in 0.9 % NaCl (saline) solution. KIC mice or control littermates received two i.p. injections of 6-OHDA (100 mg/kg at 3.5 weeks, and 250 mg/kg three days later) or vehicle solutions. For surgical sympathectomy, 3.5-week-old KIC mice or control littermates were anesthetized with 100 mg/kg ketamine and 10 mg/kg xylazine, while 100 µL of saline solution was injected subcutaneously to prevent dehydration during the procedure. Lidocain (10 mg/kg; Lurocaïne; Vétoquinol, Lure, France) was injected subcutaneously along the incision line and a midline laparotomy was performed to visualize the coeliac plexus, celiac artery, and superior mesenteric artery (AMS). Nerve bundles surrounding the AMS were severed using micro-dissection forceps under a binocular microscope (Leica, Wetzlar, Germany). After nerve sectioning, the abdominal cavity was washed with pre-warmed saline solution at 37 °C, after which abdominal muscle and skin closure was performed using 6-0 (cat. #0320543; Ethicon, Cincinnati, OH) and 4-0 vicryl sutures (cat. #0320570), respectively. Buprenorphine (0.1 mg/kg; Bupaq; Virbac, Carros, France) was injected intraperitoneally after surgery and the following day. Sympathectomized and control animals were sacrificed at 6.5 weeks of age to collect primary tumors or euthanized at the endpoint requested by the Ethics Committee for survival assays.

### Liposome preparation and injections

Liposomes encapsulating DxR were prepared and modified for CD163 targeting as previously described ^26^. KIC mice were injected with 1 mg/kg liposomes (diluted in 100 µL PBS) by retro-orbital injection of the venous sinus twice a week, with the first injection performed one day before treatment with AA or 6-OHDA. For injections, mice were anesthetized with 2 % isoflurane and a drop of ophthalmic anesthetic (1 % tetracaine; Laboratoire Tvm, Lempdes, France) was placed on the injection-receiving eye.

### Orthotopic syngeneic transplantation model

PK4A-Luc cells 24 were cultivated in DMEM GlutaMAX medium (cat. #10566016; Gibco; Thermo Fisher Scientific) supplemented with 10 % fetal bovine serum and 1 % penicillin/streptomycin (cat. #15140122; Gibco). FBV/C57BL/6 mice received 6-OHDA or vehicle solution at 3.5 weeks as described in the Materials and Methods. At 7 weeks, mice were anesthetized using ketamine (100 mg/kg) and xylazine (10 mg/kg), with 100 µL saline solution subcutaneously injected to prevent dehydration. Lidocain (10 mg/kg) was injected subcutaneously along the incision line and a midline laparotomy was performed to exteriorize the pancreatic tail. PK4A-Luc cells (150.000 cells) in 30 µL culture medium were injected into the tail of the pancreas using an insulin syringes. The abdominal cavity was washed with pre-warmed saline solution at 37 °C, after which abdominal muscle and skin closure was performed using 6-0 and 4-0 vicryl sutures, respectively. Buprenorphine (0.1 mg/kg) was injected intraperitoneally after surgery and the following day.

### Bioluminescence imaging and analysis

Bioluminescence was recorded just after transplantation of PK4A-Luc cells and every other day until day ^8^. Bioluminescence signals were induced by i.p. injection of 150 mg/kg Luciferin-EF (E6552; Promega, Madison, WI) in PBS, 10 min before in vivo imaging. Mice anesthetized with 2 % isoflurane were imaged with a Photon Imager (Biospace Lab, Nesles-la-Vallée, France). For each mouse, we plotted the logarithm of bioluminescence with respect to time (in days). Gompertz growth models were fitted to the data of each mouse. To compare sympathectomized mice (6-OHDA) with the control group (AA), a Bayesian hierarchical model was designed that includes a mouse effect on all three parameters of the parsimonious Gompertz growth model, with heteroscedasticity to take into account the possibility of a larger measurement error when bioluminescence temporarily decreases over time. Computation of the posterior distribution was performed using Stan software (New BSD Licence) with a NUTS sampling algorithm. The initial growth rate was evaluated with the first derivative at time t = 0 of the fitted curves on the logarithmic scale. Posterior distribution at the level that compares both groups was carefully analyzed, which revealed that—at a posterior probability of approximately 94 %—the initial growth rate is larger for the 6-OHDA group than for the controls. After transferring on Jeffreys’ scale to interpret the Bayesian statistical hypothesis testing, the posterior probability of 94 % indicated that the data provide “strong evidence” in support of this conclusion.

### PDX tumor models

Establishment and maintenance of PDX via successive subcutaneous transplantation have been previously described ^23^. Freshly resected xenograft (5 mm^3^) was sewn onto the pancreas of 8 week-old Swiss Nude mice (Crl:NU(Ico)-*Foxn1*nu) anesthetized with 2 % isoflurane. Mice were monitored twice a week for tumor development and sacrificed at the endpoint requested by the Ethics Committee.

### Histology and immunohistochemistry

Mice were anesthetized by i.p. injection of 100 mg/kg ketamine (Imalgene; Merial, Lyon, France) and 10 mg/kg xylazine (Rompun; Bayer, Leverkusen, Germany) and intracardially perfused with 20 mL phosphate-buffered saline (PBS) followed by 30 mL 4 % paraformaldehyde (PFA) in PBS. Dissected pancreata were post-fixed overnight at 4 °C in 4 % PFA in PBS and then rinsed three times in PBS. Hematoxylin phloxine saffron stain (HPS) and Masson’s Trichome stain (cat. #25088; Polyscience, Warrington, PA) were performed on 5 µm-thick paraffin sections. Pimonidazole staining was performed using the Hydroxyprobe^TM^-1 Omni Kit (cat. #HP3-1000Kit; Hypoxyprobe, Burlington, MA) according to manufacturer’s instructions. For immunohistochemistry, 14 µm-thick cryostat sections were washed in PBS, blocked for 1 h in blocking solution (2 % donkey normal serum, 2 % bovine serum albumin, and 0.01 % Triton X-100 in PBS) and incubated overnight at 4 ºC with primary antibodies diluted in blocking solution. After several washes in PBS, sections were incubated for 2 h with secondary antibodies. Biotinylated antibodies were detected using the Vectastain ABC peroxidase kit (cat. #PK-6100; Vector Laboratory). Antibodies used are listed in Supplementary Table S3.

### Patients and survival analysis

Patients (n = 34) included in the retrospective cohorts developed PDAC and underwent diagnostic core needle biopsy or surgical resection at North Hospital, Marseille, France. Patients were diagnosed between 2005 and 2009 and followed until March 2010. Tumors were histologically staged according to the 7th edition of the American Joint Committee on Cancer (AJCC)–Union for International Cancer Control (UICC) tumor node metastasis (TNM) classification. The study included only adenocarcinomas and histological variants, as defined by the World Health Organization (WHO) classification 2010. Data on patient demographics, diagnosis, surgical resection, and outcome were retrieved from medical records in the hospital database. The study was approved by the Regional Ethical Review Board. Written or verbal informed consent from participant patients was not obtained specifically for this study, because (1) only histopathological samples for which patients had consented to inclusion and storage in the biobank, according to biobank legislation, were included in the study; and (2) further specific written consent was considered unfeasible due to the retrospective nature of the study and the general poor prognosis associated with the tumor type in this study. Formalin-fixed, paraffin-embedded human sections were immunolabeled for CD68 and CD163 (details of antibodies are available in Supplementary Table S3) and counterstaining with Mayer’s hematoxylin (cat. #S3309; Dako, Jena, Germany). Immunohistochemical images were analyzed in collaboration with the anatomical pathology laboratory of Hospital North using Calopix software (TRIBVN, Châtillon, France) and recorded in a database. The cohort was separated into two groups according to the level of CD163 and CD68 expression in their tumors (high- and low-expression groups with 0.65 as threshold) through a procedure that maximizes differences in survival distributions. We used Kaplan–Meier survival curves, with time to death censored at the end of follow-up as the outcome.

### Statistical analysis

Z-score analysis was performed in MATLAB (MathWorks, Natick, MA). For each variable, the Z-score was defined as,

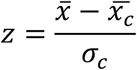

where 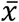 is the mean of the variable, 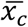 is the mean of the variable in control tissues, and *σ_c_* is the standard deviation in controls. PCA and hierarchical clustering were performed in R version 3.3.3. Other statistical analyses were performed using Graphpad Prism version 6 (GraphPad Software Inc., La Jolla, CA). Normal distribution of the data was examined using the D’Agostino–Pearson omnibus, Shapiro-Wilk, and Kolmagorov-Smirnov tests. If data were nonparametric, the Mann–Whitney test was used to compare two group means. The estimate of variance was determined by the standard deviation of each group using an F test. If the data were parametric and the variances equal, we used an unpaired *t*-test, and if the variances were different we used an unpaired *t*-test with Welch’s correction. For survival analysis, comparisons between Kaplan–Meier survival curves were performed using log-rank (Mantel–Cox) and hazard ratio tests.

## Acknowledgements

We thank Marie Falque (Institut de Biologie du Développement de Marseille, IBDM), Angélique Puget (IBDM), and Dolores Barea (Centre de Recherche en Cancérologie de Marseille) for their substantial help with the experiments, and François Michel at INMED Imaging facility (InMAGIC) for his help with LSFM. This work was supported by Centre National de la Recherche Scientifique (CNRS), France; Aix Marseille Université, France; grants from Fondation ARC (PJA 20151203159) and INSERM (Program HTE PITCHER 201609) to F.M.; grants from FRM (AJE20150633331), ANR (ANR-16-ACHN-0011) and A*midex Chaire d’excellence to SvdP; grants from INCa (PLBIO15-217), Fondation ARC (PJA 20181208127) and INSERM (BBG/2017) to F.G.; grant from the Novo Nordisk Foundation (NNF14OC0008781) to A.E.; grant from INCa, la Ligue contre le cancer and Fondation ARC (PAIR Pancreas 186738) to F.M., R.T., S.P., F.H and J-Y. S, The France-BioImaging infrastructure is supported by the Agence Nationale de la Recherche (ANR-10-INBS-04-01, ‘Investissements d’Avenir’). J.G. is a recipient of doctoral fellowships from MESRI and from Fondation ARC; A.L. is a recipient of a doctoral fellowship from MESRI; T-T-H. N. is a recipient of a Program 911-VIED doctoral fellowship.

## Author Contributions

Acquisition of data: J.G., C.D., A.L., T-T-H.N, S.C.. Analysis and interpretation of data: J.G., C.D., A.L., T-T-H.N., P.P., F.H., J-Y.S., R.T., S.C., F.M. Technical and material support: J.N., F.G., M.B., N.D., A.E., T.L., J-Y.S., S.P., R.T. Study supervision: S.C., F.M.

