## Supplementary data for "Sympathetic axonal sprouting induces changes in macrophage populations and protects against pancreatic cancer"

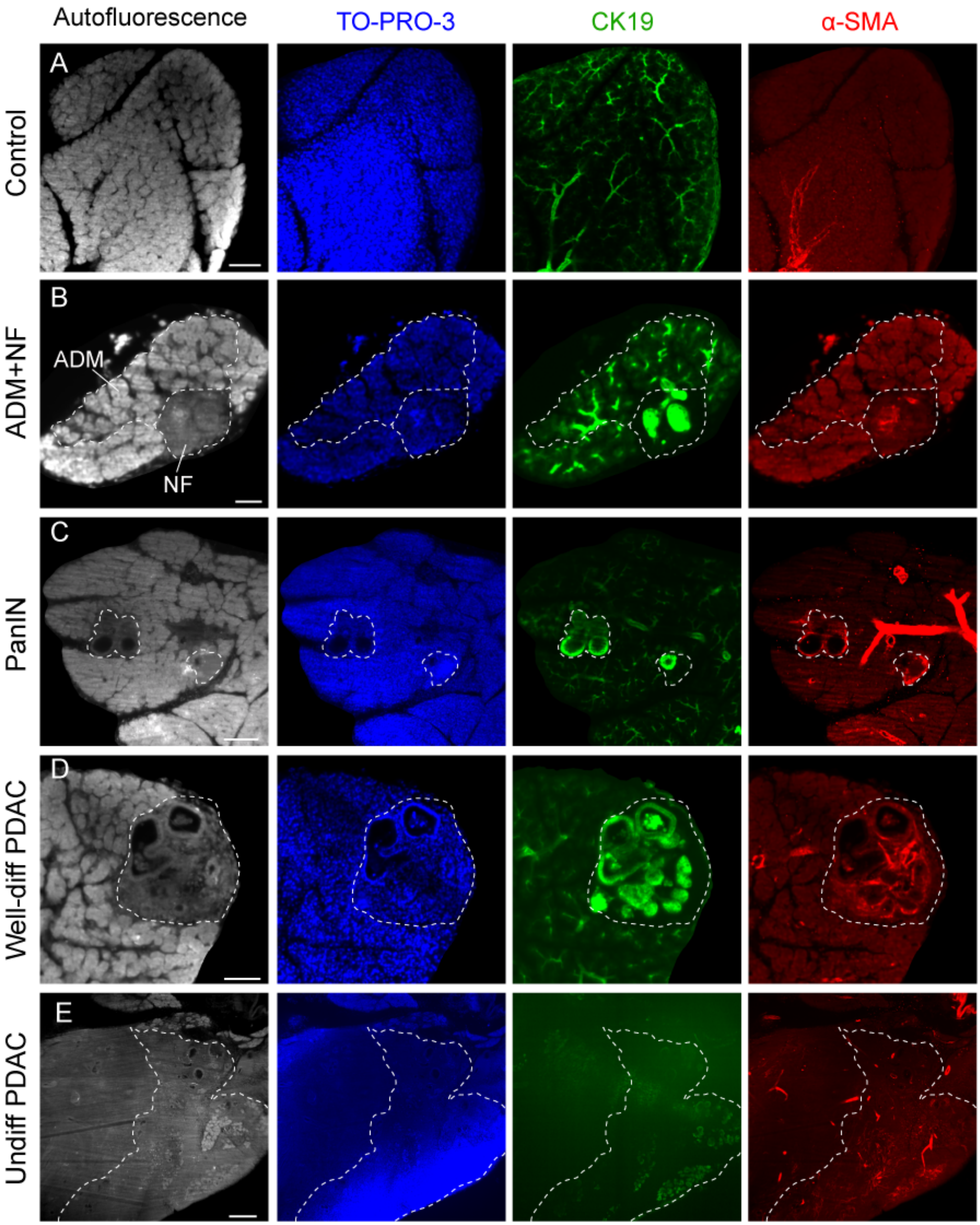

**Supplementary Figure S1. Identification of regions of interest (ROI) using tissue autofluorescence.** **A**–**E**, Optical sections through cleared pancreata of 8-week-old control (**A**) and KIC **(B–E)** mice. From left to right, panels show the acquisition of tissue autofluorescence (imaged in the blue-green spectrum with 480 nm laser excitation), nuclear counterstaining with TO-PRO-3, and immunofluorescent signals of anti-CK19 and anti-α-SMA antibodies. Images show normal acinar tissue (**A**), ADM and NF (**B**), PanIN lesions (**C**), well-differentiated PDAC (**D**), and undifferentiated/anaplastic PDAC regions (**E**). ROI are delimited by a white dashed line. In control pancreas, CK19 is expressed by epithelial cells of pancreatic ducts and α-SMA is expressed by smooth muscle cells in vessel walls. In pretumor and tumor lesions, CK19 is expressed by cancer cells and α-SMA is expressed by cancer-associated fibroblasts (CAFs). Tissue autofluorescence provides sufficient histological information to identify the different stages of disease progression. The identity of the pancreatic lesions shown above has been confirmed by an anatomopathologist. Scale bars = 50 µm (**A**, **B**), 150 µm (**C**), 100 µm (**D**), and 300 µm (**E**).

A IMAGE TREATMENT

| Imaris tools | Extracted data |
|---|---|
| **Surface** | - *Volume* of ROI |
| **Surface** | - *Volume* of duct lumen |
| **Filament tracer** | - *Dendrite Length* (sum)<br>- *Dendrite Length* (mean)<br>- *Dendrite Straightness* (mean)<br>- *Filament No. Dendrite Branch Pts*<br>- *Filament No. Sholl Intersections* |
| **Surface** | - *Volume* of blood vessels |
| **Surface-Surface Contact Area** | - *% SurfaceArea coverage*<br>- *Contact surface area*<br>- *Number of Disconnected Components*<br>- *Disconnected Components area (mean)* |

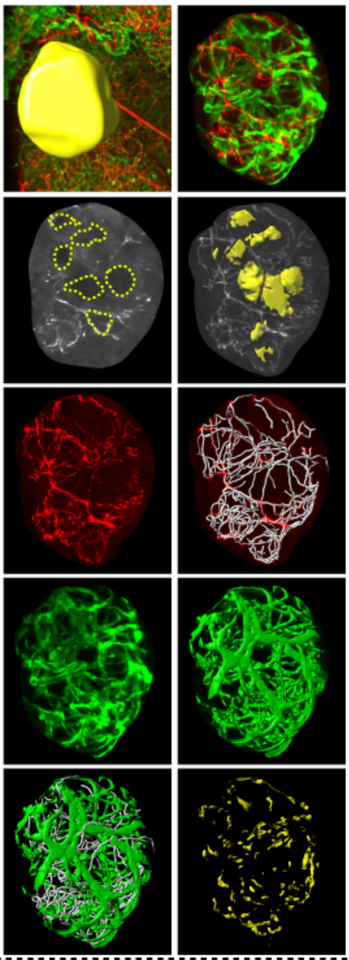

B DATA ANALYSIS

AXONAL NETWORK

| Paramater | Unit |
|---|---|
| Axon density | nm / μm³ |
| Branching | branch points / mm |
| Branch lenght | μm |
| Branch straightness | - |
| Branching pattern | - |

AXON / BV INTERACTION

| Paramater | Unit |
|---|---|
| % Contact axon / BV | % |
| Contact size | μm² |
| Contact fragmentation | contacts / μm² |
| Contact density | contacts / μm³ |
| BV density | - |
| % Contact BV / axon | % |
| Contact / Volume BV | contacts / μm³ |

C STATISTICAL ANALYSIS

→ Z-score → PCA

**Supplementary Figure S2. Summary of image treatment and data analysis.**

**A**, LSFM images were processed with Imaris software. ROI were identified via tissue autofluorescence (imaged at 488 nm excitation). A mask was created around each ROI using the “Surface” tool, after which the ROI volume was collected in the statistics tab. In non-invasive pancreatic lesions and PDAC, masks were created around the duct-like cavities using the “Surface” tool and the volume of the cavities (empty space) was collected and deduced from the ROI volume. Axonal networks were reconstructed manually using “Filament tracer.” The following parameters were collected in the statistics tab: dendrite length (sum and mean values), defined as the length of each axon branch; dendrite straightness (mean value), defined as the ratio between branch length and radial distance between two branch points; filament no. Sholl intersections, defined as the number of branch intersections per concentric spheres originating at the centroid of the ROI; and filament no. dendrite branch pts, defined as the number of branching points in the entire axon network. The “Surface” tool was used to reconstruct vascular networks and collect the volume occupied by the blood vessels. Finally, the MATLAB plug-in Imaris XTension “Surface-Surface contact area” was used to generate the contact surface areas between axonal and blood vessels networks. In the statistic tab, we collected the number of disconnected components, defined as the number of contacts between the two surfaces; the disconnected components area (mean value), defined as the area of each contact region; and the contact surface area; defined as the total surface of contacts. We then collected the % Surface Area coverage, defined as the percentage of axon surface in contact with blood vessels (% contact axon/BV) or as the percentage of blood vessel surface in contact with axons (% contact BV/axon). **B, C**, The extracted data were used to calculate 12 parameters describing the morphology of axonal networks and interactions between axons and blood vessels (BV) (**B**). Details of the calculations are provided in Supplementary Table 1. Statistical analysis was performed using MATLAB and R software (**C**).

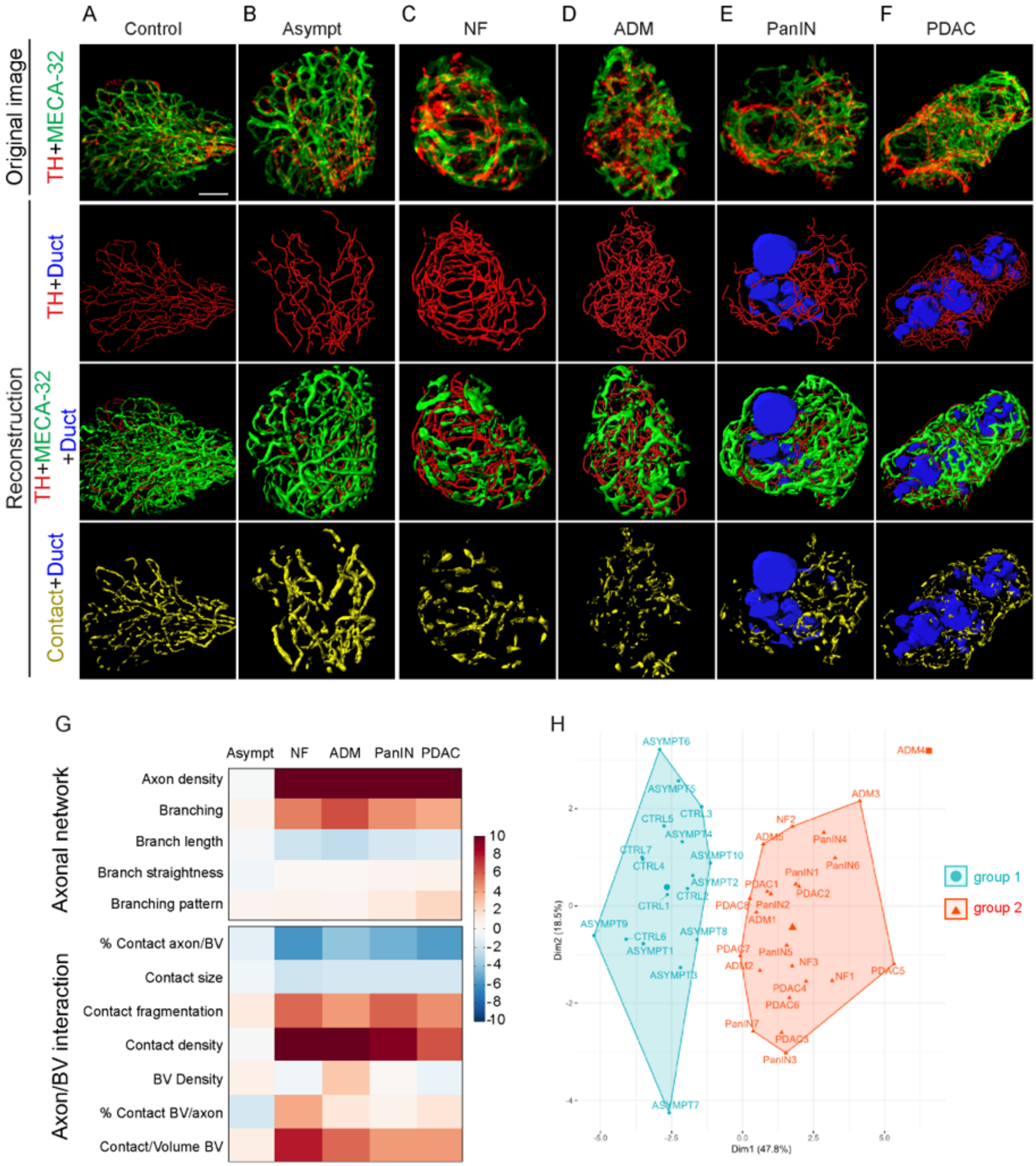
A Control
B Asympt
C NF
D ADM
E PanIN
F PDAC
Original image
TH+MECA-32
Reconstruction
TH+Duct
TH+MECA-32 +Duct
Contact+Duct
G
Asympt NF ADM PanIN PDAC
Axonal network
Axon density
Branching
Branch length
Branch straightness
Branching pattern
Axon/BV interaction
% Contact axon/BV
Contact size
Contact fragmentation
Contact density
BV Density
% Contact BV/axon
Contact/Volume BV
10 8 6 4 2 0 -2 -4 -6 -8 -10
H
group 1
group 2
Dim1 (47.8%)
Dim2 (18.5%)
ASYMPT6
ASYMPT7
ADM4
ADM3
PDAC5
PanIN3

**Supplementary Figure S3. 3D visualization and statistical analysis of sympathetic axon and blood vessel networks in KPC pancreas.**

**A–F,** Representative images of 14-week-old control (**A**) or KPC (**B–F**) pancreata immunostained with anti-TH and MECA-32 antibodies (first column). 3D reconstructions of sympathetic axons (red, second column), blood vessels (green, third column), and axon/vessel surface contacts (yellow, fourth column) in normal acinar tissue (**A**), asymptomatic acinar tissue (**B**), NF (**C**), ADM (**D**), PanIN (**E**), and a well-differentiated PDAC region (**F**). Cavities of the duct-like structures are represented in blue. **G,** Heatmap of the Z-scores calculated for each of the 12 variables describing the architecture of sympathetic axons and their relationship with blood vessels. Values in 10 asymptomatic acinar regions (Asymp), 3 NF, 5 ADM, 7 PanIN, and 8 PDAC samples of 14-week-old KPC mice (n = 3) were compared with those of 7 normal acinar regions in age-matched control mice (n = 3). **H,** Factor map of the PCA performed on 40 tissue samples and 12 variables describing sympathetic innervation. Two cluster groups were identified corresponding to control and asymptomatic tissues (group 1, blue) and pre-tumor and tumor lesions (group 2, red). Scale bars = 25 μm (**C**), 50 μm (**A**, **B** and **D–F**).

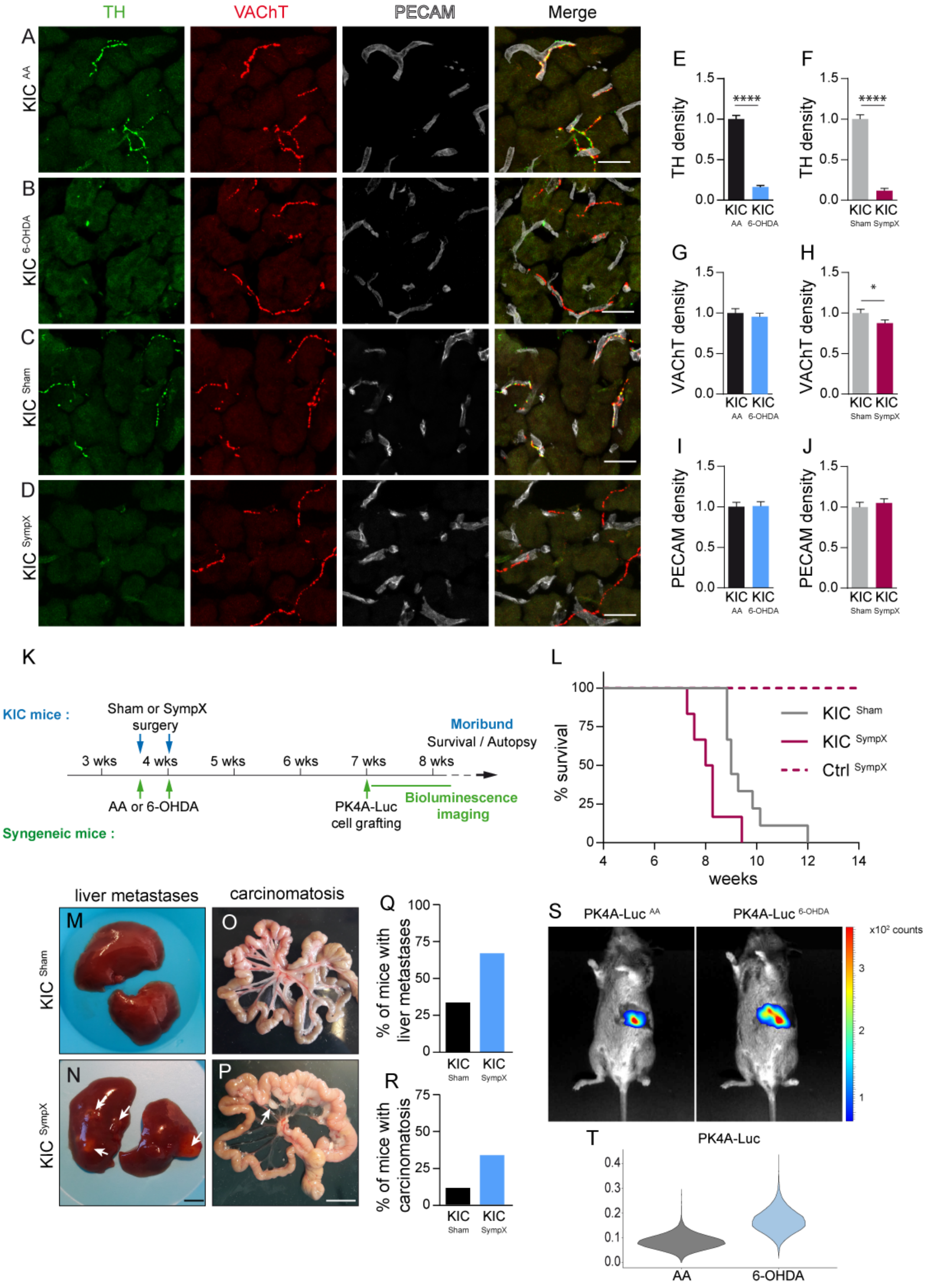
TH
VAChT
PECAM
Merge
A
KIC AA
B
KIC 6-OHDA
C
KIC Sham
D
KIC SympX
E
TH density
****
KIC KIC
AA 6-OHDA
F
TH density
****
KIC KIC
Sham SympX
G
VAChT density
KIC KIC
AA 6-OHDA
H
VAChT density
*
KIC KIC
Sham SympX
I
PECAM density
KIC KIC
AA 6-OHDA
J
PECAM density
KIC KIC
Sham SympX
K
KIC mice :
Sham or SympX surgery
3 wks
4 wks
5 wks
6 wks
7 wks
8 wks
Moribund
Survival / Autopsy
AA or 6-OHDA
PK4A-Luc cell grafting
Bioluminescence imaging
Syngeneic mice :
L
% survival
weeks
KIC Sham
KIC SympX
Ctrl SympX
liver metastases
carcinomatosis
M
N
O
P
KIC Sham
KIC SympX
Q
% of mice with liver metastases
KIC KIC
Sham SympX
R
% of mice with carcinomatosis
KIC KIC
Sham SympX
S
PK4A-Luc AA
PK4A-Luc 6-OHDA
x10² counts
T
PK4A-Luc
AA
6-OHDA

**Supplementary Figure S4. Effects of sympathectomy on tumor growth, metastatic spread, and survival.**

**A–D,** Representative images of pancreatic acinar tissue immunonostained with anti-TH, anti-VAChT, and anti-PECAM antibodies. The pancreas was collected from mice treated with AA (**A**) or 6-OHDA (**B**) or from mice that had undergone sham surgery (Sham; **C**) or surgical sympatectectomy (SympX; **D**) two weeks after chemical or surgical sympathectomy. **E–J,** Histograms representing the density of $TH^{+}$ fibers (**E, F**), $VAChT^{+}$ fibers (**G, H**), and $PECAM^{+}$ vessels (**I, J**) in the pancreas of sympathectomized mice shown in (**A-D**). Data are presented as the mean ± SEM; $p < 0.0001$ (**E**, **F**) and $p = 0.0423$ (**H**) with Mann–Whitney test. $KIC^{AA}$, n = 2 mice, 54 images; $KIC^{6\text{-}OHDA}$, n = 2 mice, 52 images; $KIC^{Sham}$, n = 2 mice, 40 images; $KIC^{SympX}$, n = 2, 36 images). **K,** Outline of the in vivo experiments. **L,** Kaplan–Meier curve comparing overall survival of KIC mice with intact pancreas ($KIC^{Sham}$, n = 9) or with surgical sympathectomy ($KIC^{SympX}$, n = 6) and control sympathectomized mice ($Ctrl^{SympX}$, n = 5). $KIC^{Sham}$ vs $KIC^{SympX}$, log-rank = 0.0109 and hazard ratio B/A = 3.179 (A = $KIC^{Sham}$ and B = $KIC^{SympX}$). **M–P,** Representative pictures of livers and intestine of $KIC^{Sham}$ and $KIC^{SympX}$ mice collected at moribund stage. **Q–R,** Graphs showing the percentage of $KIC^{Sham}$ (n = 6) and $KIC^{SympX}$ (n = 9) mice with liver metastasis (**Q**) or carcinomatosis in the intestine mesentery (**R**) at death. **S,** Bioluminescence images of AA- and 6-OHDA-treated FVB/C57BL/6 mice 8 days after orthotopic grafting of PK4A-Luc cells. **T**, Comparison of the posterior distribution of the initial growth rates of the 6-OHDA and control groups with violin plots. Numerical computations show that the 6-OHDA rate is greater than the AA rate with a posterior probability of approximately 94%. Scale bars = 50 µm (**A–D**), 5 mm (**M**, **N**), and 1 cm (**O**, **P**).

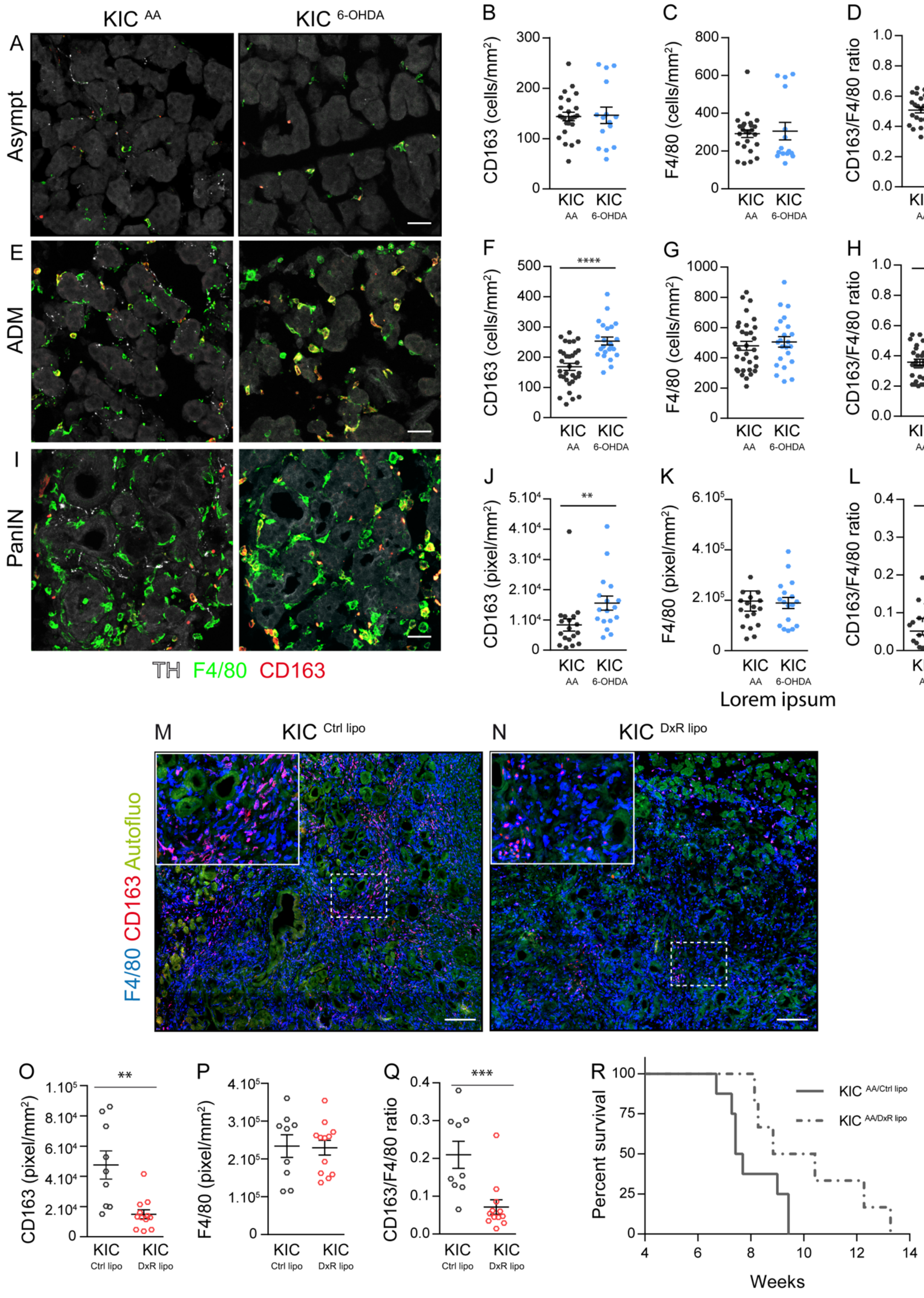

KIC AA
KIC 6-OHDA
A
Asympt
E
ADM
I
PanIN
TH F4/80 CD163
B
CD163 (cells/mm²)
C
F4/80 (cells/mm²)
D
CD163/F4/80 ratio
F
G
H
J
CD163 (pixel/mm²)
K
F4/80 (pixel/mm²)
L
Lorem ipsum
M
KIC Ctrl lipo
N
KIC DxR lipo
F4/80 CD163 Autofluo
O
P
Q
R
Percent survival
KIC AA/Ctrl lipo
KIC AA/DxR lipo
Weeks

**Supplementary Figure S5. Sympathectomy increased CD163 expression in KIC pancreas.**

**A,** Immunostaining for F4/80, CD163, and TH in asymptomatic acinar tissue (Asympt) of 6.5-week-old KIC mice treated with AA or 6-OHDA. **B–D**, Scattered dot plots depicting the density of CD163+ macrophages (**B**; p = 0.7720, Mann–Whitney test), F4/80+ macrophages (**C**; p = 0.3411, Mann–Whitney test), and CD163/F4/80 ratio (**D**; p = 0.9956, Mann–Whitney test) in asymptomatic regions of AA and 6-OHDA-treated KIC mice. $KIC^{AA}$, n = 4 mice, 25 images; $KIC^{6\text{-}OHDA}$, n = 3 mice, 15 images. **E**, Immunostaining for F4/80, CD163, and TH in ADM of 6.5-week-old KIC mice treated with AA or 6-OHDA. **F–H**, Scattered dot plots depicting the density of CD163+ macrophages (**F**; p < 0.0001, unpaired *t*-test), F4/80+ macrophages (**G**; p = 0.5667, unpaired *t*-test), and CD163/F4/80 ratio (**H**; p < 0.0001, Mann–Whitney test) in the ADM of AA- and 6-OHDA-treated KIC mice. $KIC^{AA}$, n = 6 mice, 33 images; $KIC^{6\text{-}OHDA}$, n = 4 mice, 23 images. **I**, Immunostaining for F4/80, CD163, and TH in PanIN lesions of 6.5-week-old KIC mice treated with AA or 6-OHDA. **J–L**, Scattered dot plots depicting the density of CD163+ macrophages (**J**; p < 0.0001, Mann–Whitney test), F4/80+ macrophages (**K**; p = 0.0024; Mann–Whitney test), and CD163/F4/80 ratio (**L**; p = 0.0153, Mann–Whitney test) in PanIN of AA- and 6-OHDA-treated KIC mice. $KIC^{AA}$, n = 4 mice, 18 images; $KIC^{6\text{-}OHDA}$, n = 4 mice, 17 images. **M, N,** Immunolabeling with anti-F4/80 and anti-CD163 antibodies of PDAC sections from 6.5-week-old KIC mice injected twice a week with Ctrl lipo or DxR lipo for 3 weeks. Tissue histology is revealed via autofluorescence. **O–Q,** Scattered dot plots depicting the density of CD163+ macrophages (**O**; p = 0.0013, Mann–Whitney test), F4/80+ macrophages (**P**; p = 0.9001, unpaired *t*-test), and CD163/F4/80 ratio (**Q**; p = 0.0003, Mann–Whitney test) in PDAC of 6.5-week-old KIC mice treated for 3 weeks with Ctrl lipo ($KIC^{Ctrl\ lipo}$, n = 2 mice, 9 images) or DxR lipo ($KIC^{DxR\ lipo}$, n = 3 mice, 12 images). **R**, Kaplan–Meier curve comparing overall survival of AA-treated KIC mice injected with Ctrl lipo ($KIC^{AA/Ctrl\ lipo}$, n = 8, same as in Figure 7I) or DxR lipo ($KIC^{AA/DxR\ lipo}$, n = 8). $KIC^{AA/Ctrl\ lipo}$ vs. $KIC^{AA/DxR\ lipo}$, log-rank = 0.0109 and hazard ratio B/A = 3.665 (A = $KIC^{AA/Ctrl\ lipo}$ and B = $KIC^{6\text{-}\ AA/DxR\ lipo}$). Scale bars= 50 µm (**A**, **E**, and **I**), 200 µm (**M**, **N**). Error bars represent median ± SEM.

**Supplementary Table S1: Definition of parameters describing axonal morphology and relationships with blood vessels**

| Parameters | Algorithm | Units |
|---|---|---|
| Axon density | $\frac{Dendrite\ length\ \text{(sum)}\ X\ 1000}{Volume\ \text{of the ROI} - \sum Volume\ \text{of the cavities}}$ | nm / $\mu m^3$ |
| Branching | $\frac{Filament\ No.Dendrite\ Branch\ Pts\ X\ 1000}{Dendrite\ length\ (Sum)}$ | branch points / mm |
| Branch length | *Dendrite Length* (mean) | µm |
| Branch straightness | *Dendrite Straightness* (mean) | -- |
| Branching pattern | *Filament No. Sholl Intersections* | -- |
| % contacts axon/BV | *% SurfaceArea coverage*<br>(primary surface = surface of filaments) | % |
| Contact size | *Disconnected Components area* (mean) | $\mu m^2$ |
| Contact fragmentation | $\frac{Number\ of\ Disconnected\ Components}{Contact\ surface\ area}$ | contacts /$\mu m^2$ |
| Contact density | $\frac{Number\ of\ Disconnected\ Components}{Volume\ \text{of the ROI} - \sum Volume\ \text{of the cavities}}$ | contacts/ $\mu m^3$ |
| BV density | $\frac{Volume\ \text{of the BV}}{Volume\ \text{of the ROI} - \sum Volume\ \text{of the cavities}}$ | -- |
| % contact BV/axon | % SurfaceArea coverage<br>(primary surface = vascular surface) | % |
| Contact/Volume BV | $\frac{Number\ of\ Disconnected\ Components}{Volume\ \text{of the BVs}}$ | contacts /$\mu m^3$ |

**Supplementary Table S3: Referential list of antibodies used**

| Antibody name | Host animal | Diluti on | Source | Identifier |
|---|---|---|---|---|
| **Primary antibodies** | | | | |
| Anti-Tyrosine Hydroxylase (TH) antibody | rabbit | 1:200 | Abcam | Cat # ab76442, RRID: AB_1524535 |
| Anti-Tyrosine Hydroxylase (TH) antibody | chicken | 1:200 | Aves Labs | Cat # TYH, RRID: AB_10013440 |
| Anti-Vesicular Acethylcholine Transporter (VAChT) antibody | rabbit | 1:250 | Synaptic Systems | Cat # 139 103, RRID:AB_887864 |
| Anti- CD31 Platelet Endothelial Cell Adhesion Molecule (PECAM) antibody | rat | 1:400 | BD BioScience | Cat # 553370, RRID: AB_394816 |
| Anti-Synaptophysin 1 antibody | guinea pig | 1:400 | Synaptic Systems | Cat #101 004, RRID: AB_1210382 |
| Anti-F4/80 antibody | rat | 1:50 | Santa Cruz Biotechnology | Cat # sc-71088, RRID: AB_1122714 |
| Anti-CD163 antibody | rabbit | 1:1000 | non-commercial | (1) |
| Anti-CD45 antibody | rat | 1:1000 | Thermo Fisher Scientific | Cat # 13-0451-82, RRID: AB_466446 |
| Anti-Ki-67 antibody | mouse | 1:400 | BD Biosciences | Cat # 550609, RRID: AB_393778 |
| Anti-Actin smooth muscle (□-SMA) antibody | rabbit | 1:300 | Abcam | Cat # ab5694, RRID: AB_2223021 |
| Anti-Mouse cytokeratin 19 (CK-19) antibody | rat | 1:40 | DSHB | Cat # TROMA-III, RRID: AB_2133570 |
| Anti- Panendothelial Cell Antigen Monoclonal Antibody, Clone MECA-32 | rat | 1:10 | BD Biosciences | Cat # 550563, RRID: AB_393754 |
| Anti-Insulin antibody | rabbit | 1:200 | Cell Signaling Technology | Cat # 4590, RRID : AB_659820 |
| Anti-Doublecortin (DCX) antibody | rabbit | 1:500 | Abcam | Cat # ab18723, RRID: AB_732011 |
| Anti-CD163 (M96) antibody | rabbit | 1:100 | Santa Cruz Biotechnology | Cat # sc-33560, RRID : AB_2074556 |
| **Secondary antibodies** | | | | |
| Alexa Fluor 568-conjugated anti-Chicken IgY (H+L) antibody | goat | 1:500 | Thermo Fisher Scientific | Cat# A-11041, RRID:AB_2534098 |
| Cy3- conjugated anti-Guinea Pig IgG (H+L) antibody | donkey | 1:500 | Jackson Immuno Research Labs | Cat# 706-165-148, RRID:AB_2340460 |
| Alexa Fluor 488-conjugated anti-Rabbit IgG (H+L) antibody | donkey | 1:500 | Thermo Fisher Scientific | Cat# A-21206, RRID:AB_2535792 |
| Alexa Fluor 568-conjugated anti-Rabbit IgG (H+L) antibody | donkey | 1:500 | Thermo Fisher Scientific | Cat#A10042, RRID: AB_2534017 |
| Alexa Fluor 647- conjugated anti-Rabbit IgG (H+L) antibody | donkey | 1:500 | Thermo Fisher Scientific | Cat# A-31573, RRID: AB_2536183 |

| | | | | |
|---|---|---|---|---|
| Alexa Fluor 790-conjugated anti-Rabbit IgG (H+L) antibody | donkey | 1:500 | Jackson Immuno Research Labs | Cat# 711-655-152, RRID: AB_2340628 |
| Cy3-conjugated anti-Rabbit IgG (H+L) antibody | donkey | 1:500 | Jackson Immuno Research Labs | Cat# 711-165-152, RRID: AB_2307443 |
| Alexa Fluor 647-conjugated anti-Rat IgG (H+L) antibody | donkey | 1:500 | Jackson Immuno Research Labs | Cat#712-605-153, RRID: AB_2340694 |
| Alexa Fluor 790-conjugated anti-Rat IgG (H+L) antibody | donkey | 1:500 | Jackson Immuno Research Labs | Cat# 712-655-153, RRID: AB_2340701 |
| Cy3-conjugated anti-Rat IgG (H+L) antibody | donkey | 1:500 | Jackson Immuno Research Labs | Cat# 712-165-153, RRID: AB_2340667 |

1. Etzerodt A, Moestrup SK. CD163 and inflammation: Biological, diagnostic, and therapeutic aspects. Antioxidants Redox Signal. 2013. page 2352–63.
